# Single-molecule analysis reveals cooperative stimulation of Rad51 filament nucleation and growth by mediator proteins

**DOI:** 10.1101/2020.11.05.369629

**Authors:** Ondrej Belan, Consuelo Barroso, Artur Kaczmarczyk, Roopesh Anand, Stefania Federico, Nicola O’Reilly, Matthew D. Newton, Erik Maeots, Radoslav I. Enchev, Enrique Martinez-Perez, David S. Rueda, Simon J. Boulton

## Abstract

Homologous recombination (HR) is an essential DNA double-strand break (DSBs) repair mechanism frequently inactivated in cancer. During HR, RAD51 forms nucleoprotein filaments on RPA-coated resected DNA and catalyses strand invasion into homologous duplex DNA. How RAD51 displaces RPA and assembles into long HR-proficient filaments remains uncertain. Here, we employ single-molecule imaging to investigate the mechanism of nematode RAD-51 filament growth in the presence of BRC-2 (BRCA2) and RAD-51 paralogs, RFS-1/RIP-1. BRC-2 nucleates RAD-51 on RPA-coated DNA, while RFS-1/RIP-1 acts as a ‘chaperone’ to promote 3’ to 5’ filament growth via highly dynamic engagement with 5’ filament ends. Inhibiting ATPase or mutation in RFS-1 Walker box leads to RFS-1/RIP-1 retention on RAD-51 filaments and hinders growth. *rfs-1* Walker box mutants display sensitivity to DNA damage and accumulate RAD-51 complexes non-functional for HR *in vivo*. Our work reveals the mechanism of RAD-51 nucleation and filament growth in the presence of recombination mediators.

## INTRODUCTION

Homologous recombination (HR) is a largely error-free mechanism of DNA double-strand break (DSB) repair. DSBs can arise spontaneously as a result of replication fork stalling or collapse or following exposure to DNA damaging agents, such as ionising radiation. HR repair is also essential to produce inter-homolog crossovers necessary for correct chromosome segregation at the first meiotic division (Chapman et al., 2012).

HR is a complex process consisting of several conserved steps. First, broken double-stranded DNA (dsDNA) ends are nucleolytically processed in a 5’→3’ direction, yielding 2-4 kb of single-stranded DNA (ssDNA) coated by Replication Protein A (RPA). Strongly-bound RPA is displaced by Rad51, an eukaryotic recombinase that forms helical nucleoprotein filaments with ssDNA. Within the filament, DNA is extended 1.5-fold over the dsDNA contour length, with a stoichiometry of 3 nt per RAD51 monomer and 6 RAD51 monomers per helical turn (Xu et al., 2017). Rad51 filaments then search for homologous DNA sequence within sister chromatids or homologous chromosomes, followed by strand invasion and displacement loop (D-loop) intermediate formation. The invading 3’ DNA end is extended by polymerases and processed by multiple redundant pathways to complete repair and restore the broken DNA strand (Chapman et al., 2012).

Given the lower affinity of Rad51 for ssDNA, compared to RPA, mediator proteins that promote RPA displacement are crucial for efficient HR. In higher eukaryotes, BRCA2 and Rad51 paralogs (RAD51B, RAD51C, RAD51D, XRCC2 and XRCC3 in human cells) are amongst the most critical mediator proteins. Loss of BRCA2 or individual RAD51 paralogs results in sensitivity to DSB-inducing agents, loss of Rad51 foci at sites of DSBs in cells and defective HR repair (Chapman et al., 2012). Mutations in BRCA2 or Rad51 paralogs confer hereditary breast, ovarian and other cancers (King et al., 2003; Loveday et al., 2011; Meindl et al., 2010).

Although BRCA2 and Rad51 paralogs are required for efficient Rad51 focus formation/stabilisation *in vivo*, a clear mechanistic understanding of how this is achieved is lacking. Biochemical studies have shown that sub-stoichiometric amounts of full-length human BRCA2 promote nucleation of RAD51 on ssDNA and enhance its strand exchange activity in bulk assays (Jensen et al., 2010; Liu et al., 2010; Thorslund et al., 2010). BRCA2 also binds RAD51-ssDNA filaments and inhibits Rad51 ATPase activity, thereby suppressing Rad51 release from DNA (Jensen et al., 2010; Petalcorin et al., 2007). While nematode Rad51 paralogs (RFS-1/RIP-1) do not bind to free RAD-51 in solution, they can bind to and stabilize Rad51 filaments (Taylor et al., 2015). Furthermore, bulk studies have shown that the DNA strand exchange activity of Rad51 is stimulated by the addition of sub-stoichiometric amounts of human RAD51B-RAD51C (Sigurdsson et al., 2001) or nematode Rad51 paralog complex (Taylor et al., 2015). Mechanistically, Rad51 paralogs have been suggested to intercalate into and stabilize Rad51 filaments. Intercalated Rad51 paralogs were proposed to serve as roadblocks to prevent filament disassembly by anti-recombinases, such as Srs2 (Liu et al., 2011).

While current evidence implicates BRCA2 and Rad51 paralogs in positively regulating Rad51 function, an understanding of their dynamics during the HR reaction and the mechanisms that act to promote the growth of HR-proficient Rad51 filaments remain unclear. Single-molecule studies of the *E. coli* RecA recombinase have revealed that nucleoprotein filaments rapidly form in the presence of bacterial single-stranded binding protein (SSB) by a two-step mechanism: rate-limiting nucleation followed by rapid bi-directional filament growth with two-fold kinetic preference for the 5’→3’ direction along a ssDNA backbone (Bell et al., 2012; Galletto et al., 2006). In contrast, human Rad51 filaments formed in the presence of RPA *in vitro* are rare and grow very slowly (Candelli et al., 2014; Hilario et al., 2009). Currently, the mechanisms that promote presynaptic Rad51 filament assembly in eukaryotes remains unknown.

Here, we report a single-molecule system to monitor the real-time dynamics of nematode RAD-51-ssDNA filament assembly and how this is modulated by the recombination mediators BRC-2 and RFS-1/RIP-1. Through a combination of microfluidics, optical tweezers and fluorescence microscopy, we show that BRC-2 and RFS-1/RIP-1 act together to stimulate RAD-51 presynaptic nucleoprotein filament assembly. While BRC-2 acts primarily as a RAD-51 nucleation factor on RPA-coated ssDNA, RFS-1/RIP-1 acts on nucleated RAD51-ssDNA complexes to stimulate filament growth. This division of labour between mediator proteins allows for synergistic stimulation of presynaptic RAD-51 filament assembly. Direct real-time imaging of RFS-1/RIP-1 also revealed an unexpected and highly dynamic engagement with 5’ RAD-51 filament ends that requires ATP turnover by RFS-1. Transient RFS-1/RIP-1 binding to the nascent 5’ RAD-51 filament end prevents dissociation of RAD-51 protomers and stimulates filament growth in a 3’→5’ direction. However, blocking ATPase or mutation of the RFS-1 Walker A box increases the dwell-time of RFS-1/RIP-1 on RAD-51 filament ends, which stabilises RAD-51 on DNA, but inhibits filament growth. Finally, unlike nematode strains lacking *rfs-1*, which are sensitive to DNA damage and fail to form RAD-51 foci, *rfs-1* Walker box mutants accumulate non-functional RAD-51 foci, which form in a BRC-2 dependent manner. We propose that distinct mechanisms of the two mediator proteins act sequentially to synergistically promote efficient RAD-51 nucleation, filament growth and HR stimulation.

## RESULTS

### Differential action of mediator proteins on RAD-51 presynaptic complex assembly

To examine how recombination mediators impact on the nucleation and/or growth of Rad51 filaments in the presence of its physiological competitor RPA, we have reconstituted RAD-51 filament assembly at a single-molecule level using fluorescently labelled *C. elegans* proteins and a combination of optical tweezers, confocal fluorescence microscopy and microfluidics (C-trap set-up, Fig. 1A) (Gutierrez-Escribano et al., 2019; Newton et al., 2019). To generate an HR substrate, 48.5 kb doubly-biotinylated bacteriophage λ dsDNA was trapped between two polystyrene streptavidin-coated beads, force-melted *in situ* to produce ssDNA (Candelli et al., 2013) (Fig. S1) and then coated with RPA-eGFP fusion protein (Fig. 1A). Since the small subunit of *C. elegans* RPA remains unknown (Kim et al., 2005), which precludes the purification of a nematode RPA heterotrimeric complex, all eGFP-RPA displacement assays were performed with human eGFP-RPA. The substitution is justified given the lack of interaction between RPA and BRCA2 or RAD51 in solution (Jensen et al., 2010) and the established notion that RPA serves only as a competitor of RAD51 and can be replaced by bacterial SSB in bulk mediator assays (Jensen et al., 2010). Both BRCA2 (Jensen et al., 2010) and human RAD51 paralog complex (Sigurdsson et al., 2001) are also capable of stimulating strand exchange activity of RAD51 even in the absence of RPA under sub-saturating conditions.

**Figure 1.**
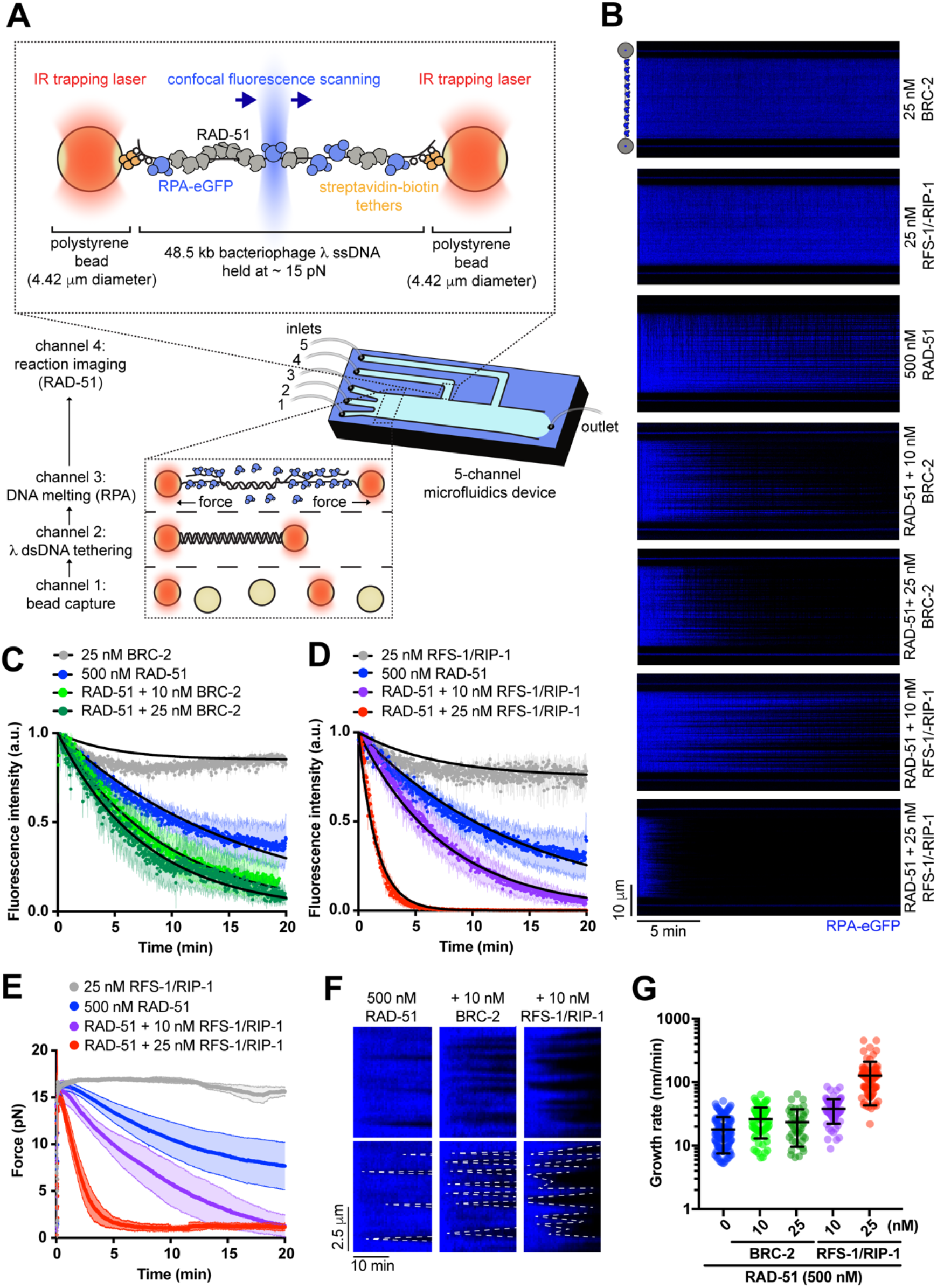
BRC-2 and RFS-1/RIP-1 display differential roles in RAD-51 filament assembly and growth on RPA-coated ssDNA. **(A)** Schematic of the experimental C-trap set-up. **(B)** Kymograph showing the displacement of RPA-eGFP by 500 nM RAD-51 in the presence or absence of BRC-2 or RFS-1/RIP-1. **(C)** Normalized fluorescence intensity for RPA–eGFP signal over time in the presence of RAD-51 and indicated amounts of BRC-2; shaded area represents SEM. (n = 3-8). Black lines represent exponential fits. **(D)** Normalized fluorescence intensity for RPA–eGFP signal over time in the presence of RAD-51 and indicated amounts of RFS-1/RIP-1; shaded area represents SEM. (n = 3-6). Black lines represent exponential fits. **(E)** Force measured between the traps as a function of time in indicated RFS-1/RIP-1 concentrations; shaded area represents SEM. (n = 3-8). **(F)** Examples of individual growing RAD-51 filaments (dark). Growth rate was measured as a slope of the border of RPA-eGFP displaced signal. **(G)** Quantification of growth rates in indicated conditions.

RAD-51 assembly was initiated by moving the traps to protein channels containing RAD-51 and/or mediator proteins (Fig. 1A). RAD-51 assembly and RPA-eGFP displacement was monitored by loss of eGFP fluorescence and simultaneous decrease in the force exerted on ssDNA that accompanies recombinase filament formation (Hegner et al., 1999).

To characterize the role of mediator proteins in our system, we performed RPA-eGFP displacement experiments with RAD-51 in the presence or absence of BRC-2, a recombination mediator and orthologue of the breast and ovarian cancer tumour suppressor protein, BRCA2 (Martin et al., 2005; Petalcorin et al., 2007; Petalcorin et al., 2006) (Fig. 1B). While RAD-51 alone slowly assembles on RPA-coated ssDNA (half-time of 10.16 ± 0.8 min, 95% CI), addition of sub-stoichiometric concentrations of BRC-2 stimulates the overall assembly rate, reducing assembly half-time to 5.3 ± 0.3 min (95% CI; Fig. 1C). We next performed RPA-eGFP displacement assays with RAD-51 in the presence of the RFS-1/RIP-1 complex. Addition of sub-stoichiometric concentrations of RFS-1/RIP-1 greatly stimulates the RAD-51 filament assembly rate, reducing assembly half-time from ∼10 min to ∼1 min (Fig. 1B, D). This is accompanied by a decrease in force measured between the optical traps from ∼15 pN to ∼1 pN (Fig. 1E), which is in agreement with increased stiffness of RAD-51-coated ssDNA (Hegner et al., 1999). Consistent with its weak ssDNA affinity (Taylor et al., 2015), RFS-1/RIP-1 did not induce any detectable RPA displacement on its own (Fig. 1B, D, E).

In addition to estimating overall assembly rate, this approach allowed us to resolve individual growing RAD-51 nuclei and measure growth rates of RAD-51 filaments (Fig. 1F, G), which confirmed that RAD-51 grows slowly (mean 17.9 ± 1.8 nm/min, 95% CI; which corresponds roughly to 38.6 ± 3.9 nt/min; 500 nM protein) compared to bacterial RecA (∼40 nm/min under similar conditions (Bell et al., 2012)). RFS-1/RIP-1 strongly stimulates growth rates of individual RAD-51 nuclei (mean of 38.1 ± 4.0 nm/min, 95% CI; which corresponds to 82.1 ± 8.6 nt/min; 10 nM RFS-1/RIP-1; Fig. 1F, G), Notably, BRC-2 had a significant, albeit modest effect on RAD-51 growth when compared to RFS-1/RIP-1 (mean of 26.4 ± 3.2 nm/min, 95% CI; which corresponds to 56.9 ± 6.9 nt/min; 10 nM BRC-2). Our data suggests that BRC-2 acts primarily as a RAD-51 nucleation factor (Shahid et al., 2014), while RFS-1/RIP-1 acts primarily as a RAD-51 filament growth factor on RPA-coated ssDNA.

### BRC-2 and RFS-1/RIP-1 synergize to ensure efficient RAD-51 presynaptic filament assembly

These distinct modes of action raised the possibility that the mediator proteins may cooperate to enhance RAD-51 filament assembly when combined in a single reaction. Inclusion of both proteins in the reaction (Fig. 2A), results in an increase in growth rate of individual RAD-51 filaments (mean of 43.8 ± 7.0 nm/min, 95% CI; which corresponds to 94.4 ± 15.1 nt/min; 10 nM BRC-2, 10 nM RFS-1/RIP-1), which was not significantly different from the RAD-51 filament growth rates measured in the presence of RFS-1/RIP-1 only (Fig. 2A, B). These results indicate that RFS-1/RIP-1 and BRC-2 do not synergize to promote RAD-51 filament growth, but rather that RFS-1/RIP-1 is the major filament growth stimulatory factor in the combined assembly reaction. However, when combined, BRC-2 and RFS-1/RIP-1 display a synergistic effect on overall RAD-51 assembly on RPA-coated ssDNA (Fig. 2A, C, D) strongly reducing overall assembly half-time to 2.4 min (95% CI). Force measurements confirmed assembly kinetics obtained from fluorescence intensity quantification (Fig. 2E). Taken together, these results further strengthen the notion that BRC-2 primarily nucleates RAD-51 on RPA-coated ssDNA, which is then extended into nascent filaments by RFS-1/RIP-1 stimulation.

**Figure 2.**
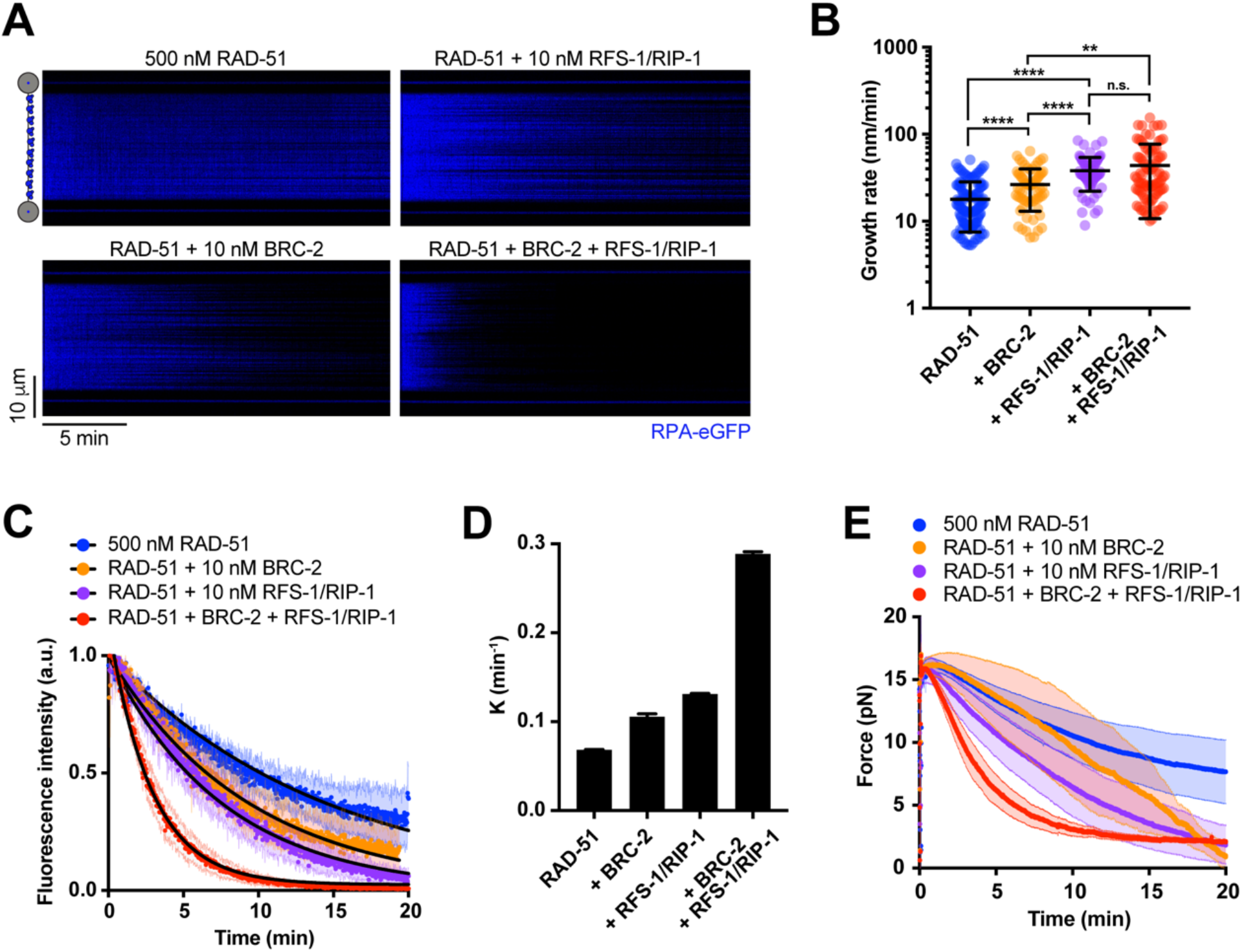
BRC-2 and RFS-1/RIP-1 synergize to ensure efficient RAD-51 filament assembly. **(A)** Kymograph showing the displacement of RPA-eGFP by 500 nM RAD-51 in the presence or absence of BRC-2 and/or RFS-1/RIP-1. **(B)** Quantification of growth rates in indicated conditions. Error bars represent SD. P > 0.05 (n.s.), P ≤ 0.05 (*), P ≤ 0.01 (**), P ≤ 0.001 (***), P ≤ 0.0001 (****). Mann-Whitney test. **(C)** Normalized fluorescence intensity for RPA–eGFP signal in the presence or absence of BRC-2 and/or RFS-1/RIP-1; shaded area represents SEM. (n = 4-8). Black lines represent exponential fits. **(D)** k_off_ values for RPA-eGFP displacement traces calculated from exponential fits to the data in 2C; error bars represent upper and lower k value limit. **(E)** Force measured between the traps as a function of time in indicated conditions; shaded area represents SEM. (n = 3-8).

### RFS-1/RIP-1 promotes filament growth in 3’ to 5’ direction

According to the orientation of the ssDNA backbone bound within the RAD51 filament, structurally distinct 3’ and 5’ filament ends are defined. Using bulk fluorescence to estimate kinetics of protein binding/dissociation from short Cy3-labelled oligonucleotides, we previously observed that RFS-1/RIP-1 binds and stabilizes the 5’ end of pre-assembled RAD-51 filaments (Taylor et al., 2016). In contrast, we did not detect any stabilization effect at the 3’ filament end. Given this previously described polarity, and RPA-eGFP displacement slopes specifically steeper in one direction observed in kymographs in the presence of RFS-1/RIP-1 compared to the conditions where RFS-1/RIP-1 was absent (Fig. 1F), we reasoned that growth stimulation promoted by RFS-1/RIP-1 might lead to preferential protomer addition at only one end of RAD-51 filaments.

To examine the polarity of filament growth, we developed a protocol to generate an asymmetrically positioned ssDNA gap of defined length on λ DNA using Cas9 nickase and DNA melting *in situ* (Fig. 3A, Fig. S2A). We confirmed the position of the ssDNA gap using eGFP-RPA fluorescence to mark the ssDNA portion of the gapped molecule (Fig. 3B, C) and applied worm-like chain model fitting to individual force-extension curves to calculate the length of ssDNA within the substrate (Fig. 3D, E). To suppress extensive RAD-51 nucleation and better resolve individual RAD-51 filaments, free RPA-eGFP was added to the imaging channel. This setup allowed us to accurately assess directionality of RAD-51 filament growth. Surprisingly, unlike RecA, which grows preferentially in a 5’→3’ direction, nematode RAD-51 displays mostly symmetric growth with only slight bias towards the 5’→3’ direction (Fig. 3F, Fig. S2B). Addition of RFS-1/RIP-1 resulted in two-fold stimulation of 3’→5’ growth rates resulting in net growth from the 5’ filament end (Fig. 3G, Fig. S2B). These results indicate that RFS-1/RIP-1-mediated growth stimulation of RAD-51 filaments is asymmetric and occurs at 5’ filament ends.

**Figure 3.**
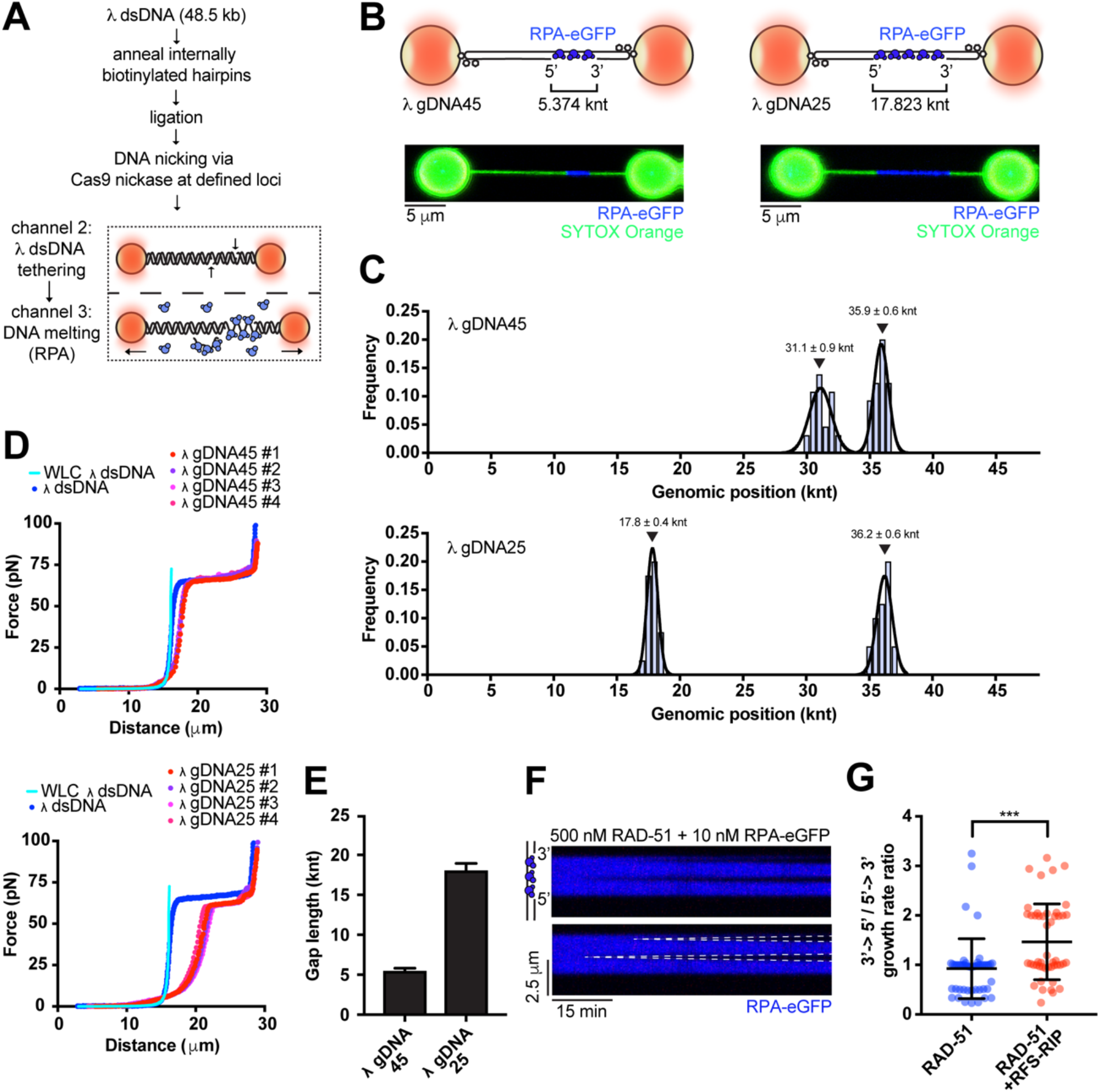
RFS-1/RIP-1 promotes filament growth in 3’ to 5’ direction. **(A)** Schematic of protocol designed to generate gapped λ DNA (gDNA) substrates. **(B)** Schematic of the two λ gDNA substrates employed to investigate RAD-51 filament growth polarity. Representative image of asymmetrically positioned RPA-eGFP coated 5 knt (λ gDNA45) and 17 knt (λ gDNA25) ssDNA gap within λ DNA held at 15 pN force. dsDNA stained by 50 nM SYTOX Orange. **(C)** Genomic position analysis of the ssDNA gap evaluated from RPA-eGFP signal boundaries. 20-32 scans analysed per each histogram. Black lines represent gaussian fits. Error represents SD. **(D)** Force-extension curves of λ dsDNA (blue) and multiple λ gDNA45 or λ gDNA25 molecules. Light blue line represents worm-like chain model (WLC) fit for 48.5 kb long dsDNA. **(E)** ssDNA gap length of λ gDNA45 and λ gDNA25 calculated from WLC model fits to traces presented in Fig. 3D. Error bars represents SD. **(F)** Examples of individual growing RAD-51 filaments (dark) on gapped DNA construct. Growth rate was measured as a slope of the border of RPA-eGFP displaced signal. **(G)** Quantification of growth rate polarity of 500 nM RAD-51 in the presence or absence of 10 nM RFS-1/RIP-1.

### RFS-1/RIP-1 acts as a molecular chaperone to stabilize RAD-51

Rad51/RecA-ssDNA filaments are in a state of dynamic equilibrium, where they grow and shrink exclusively from filament ends (Joo et al., 2006; van Mameren et al., 2009). Previous work established that human Rad51-dsDNA filaments dissociate from the ends in bursts of multiple protomers interspersed by pauses (van Mameren et al., 2009). Similar behaviour was later observed for Rad51-ssDNA complexes (Candelli, 2013). On the contrary, growth of RecA (Bell et al., 2012) and Rad51 (Candelli et al., 2014) filaments occurs predominantly by rate-limiting slow addition of monomers. Given the resolution of our setup, we were not able to accurately monitor RAD51 protomer association and dissociation events from filament ends. To circumvent this issue, we employed a previously described reverse setup (Candelli et al., 2014), where low concentrations of labelled protein is used to obtain sparse nucleation events containing only a small number of RAD-51 protomers. Single step photobleaching calibration was used to quantify the number of fluorophores present in the clusters. Given that recombinase filaments can grow and dissociate only from filament ends (Joo et al., 2006; van Mameren et al., 2009), this system allows us to assess growth and dissociation dynamics of small RAD-51 clusters and use it as a proxy for events occurring at the ends of long ssDNA RAD-51 filaments.

To investigate the molecular mechanism of RFS-1/RIP-1-mediated RAD-51 filament growth stimulation at a single-protomer level, we stoichiometrically labelled RAD-51 with fluorescein (6-FAM) (Amitani et al., 2010) (referred to as RAD-51^f^; Fig. S3A), which retained wild type levels of DNA binding (Fig. S3B) and D-loop formation activity (Fig. S3C). To visualize individual fluorescently labelled RAD-51^f^ clusters (a DNA-bound species containing 1 or more RAD-51^f^ protomers), we employed a ‘dipping protocol’ (Fig. S4A), where λ ssDNA trapped between two beads is incubated for a defined period of time in a channel containing RAD-51^f^. ssDNA bound by RAD-51^f^ is subsequently moved to a protein-free observation channel and visualized by fluorescence microscopy. Repeated incubation-detection cycles allow for kinetic analysis of RAD-51^f^ nucleation and growth. Nucleation frequency of RAD-51^f^ clusters (k_obs_) assessed after a single round of dipping displays power dependence with respect to RAD-51^f^ concentration, according to k_obs_=J[RAD-51]^n^, where J is a rate constant and n is the number of protomers in a minimal nucleation unit. In agreement with studies of bacterial RecA (n = 1.5 in ATP and n = 2.2 in ATP-γ-S; (Bell et al., 2012) and human RAD51 (n = 1.5; (Candelli et al., 2014), n = 1.6 ± 0.2 for *C. elegans* RAD-51 (Fig. S4B, C) indicating that RAD-51^f^ nucleates as a small species on ssDNA, most likely a dimer or a monomer. Consistent with our previous result (Fig. 1B, C), inclusion of BRC-2 dramatically increases nucleation frequencies in the presence of RPA in this assay (Fig. S4C). Addition of unlabelled RFS-1/RIP-1 also increased RAD-51 nucleation rates, albeit to a lesser extent than in the presence of BRC-2.

To examine the dynamics of RFS-1/RIP-1 during RAD-51 filament assembly, we fused the C-terminus of RIP-1 to a ybbr tag (Yin et al., 2006) and labelled the corresponding complex with Alexa 647 dye (referred to as RFS-1/RIP-1(A647); Fig. S5A, B). RFS-1/RIP-1(A647) retains its ability to bind RAD-51 filaments (Fig. S5C) and stimulate strand exchange (Fig. S5D). Time-resolved experiments revealed a significant accumulation of RAD-51^f^ clusters over time in the presence of RFS-1/RIP-1(A647) (Fig. 4A, B). Without RFS-1/RIP-1(A647), RAD-51^f^ clusters are highly dynamic. They bind and dissociate rapidly from ssDNA with short dwell-times (Fig. 4C, E, F). These observations are reminiscent of burst-like dissociation events reported previously at human Rad51 filament ends (van Mameren et al., 2009). Inclusion of RFS-1/RIP-1(A647) in the reaction results in a significant increase in the dwell-times of RAD-51^f^ clusters on ssDNA indicating a stabilization effect mediated by RFS-1/RIP-1 (Fig. 4D, E, F).

**Figure 4.**
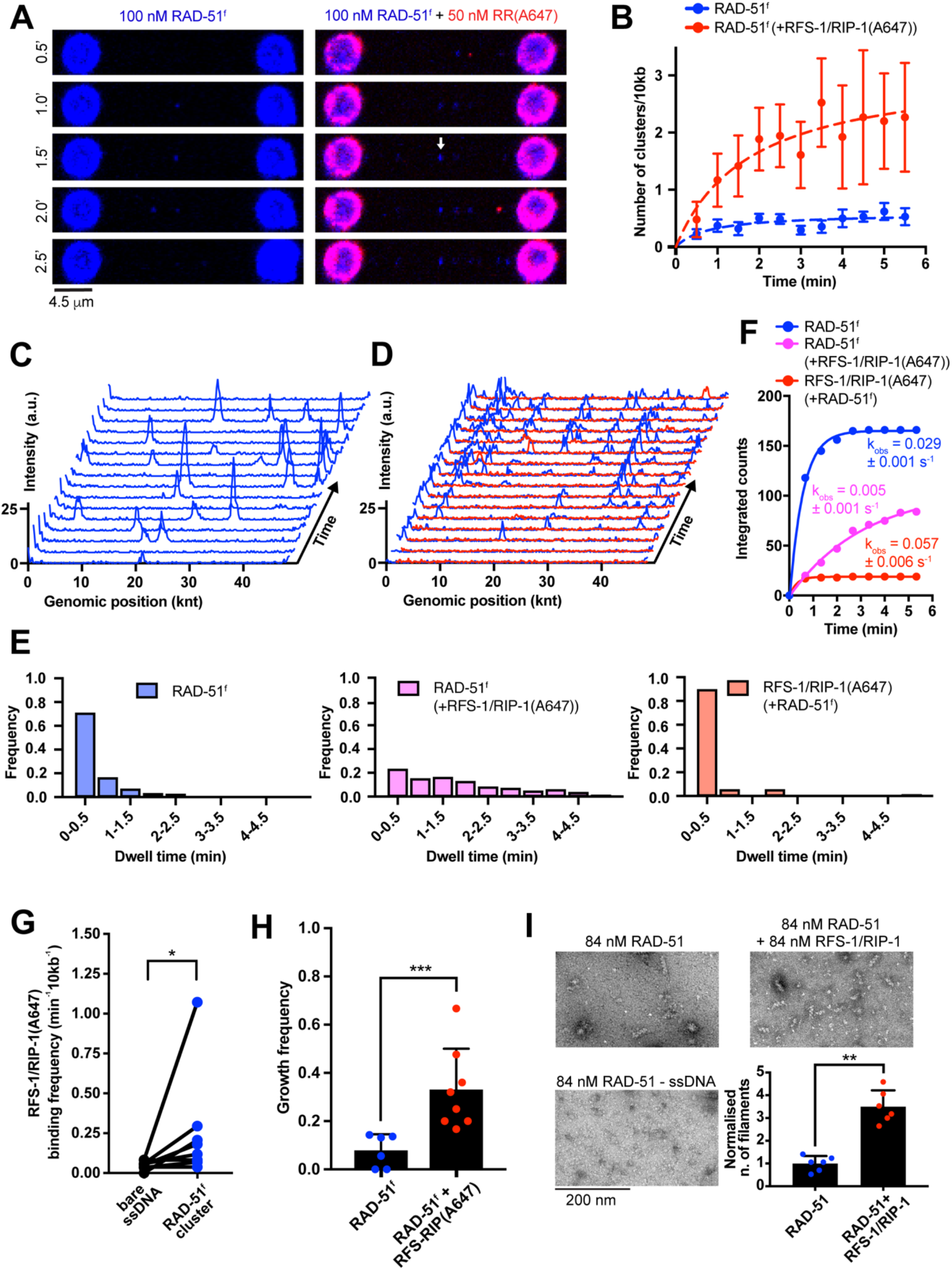
Highly dynamic RFS-1/RIP-1 complex ‘chaperones’ DNA-bound RAD-51 clusters by preventing RAD-51 dissociation. **(A)** Fluorescence images taken after subsequent RAD-51^f^ or RAD-51^f^+RFS-1/RIP-1(A647) incubation-detection cycles with the same ssDNA (no RPA) construct; cumulative incubation time (min) is indicated. 4.5 μm scale bar. Arrow indicates a growth event. RAD-51^f^ signal shown in blue. RFS-1/RIP-1(A647) signal shown in red. **(B)** Quantification of RAD-51^f^ nucleation frequency over time in the presence (n = 9) or absence (n = 11) of RFS-1/RIP-1(A647). Exponential fits are displayed. Error bars represent SEM. **(C)** Representative histogram of time-binned intensity verses genomic position on lambda DNA for RAD-51^f^ signal (blue) in the absence of RFS-1/RIP-1(A647). Each line represents 30 s timepoint. **(D)** Representative histogram of time-binned intensity versus genomic position on lambda DNA for RAD-51^f^ signal (blue) in the presence of RFS-1/RIP-1(A647) (red). Each line represents 30 s timepoint. **(E)** Histograms of dwell times of RAD-51^f^ in the absence (left panel) or presence of RFS-1/RIP-1(A647) (middle panel) or dwell times of RFS-1/RIP-1(A647) in the presence of RAD-51^f^ (right panel). Lines represent exponential fits. (upper panel: RAD-51^f^, τ ∼21.12 s, R^2^ = 0.99, n = 167; middle panel: RAD-51^f^+ RFS-1/RIP-1(A647), τ ∼150.6 s, R^2^= 0.95, n = 87; bottom panel: RFS-1/RIP-1(A647), τ ∼10.28, R^2^ = 0.99, n = 19) **(F)** Cumulative survival plots of data presented in 2e. Lines represent exponential fits. **(G)** Quantification of the frequency of RFS-1/RIP-1(A647) binding to ssDNA and RAD-51 clusters. Individual DNA molecules were analyzed and the fraction of RFS-1/RIP-1(A647) bound to ssDNA or RAD-51^f^ cluster was calculated. Each data point represents one ssDNA molecule analyzed. p = 0.01. Wilcoxon test. **(H)** Growth frequencies of RAD-51^f^ clusters (fraction of clusters on a given ssDNA molecule displaying at least one growth event – defined by an increase of protomer number by a minimum of 1) in the presence or absence of RFS-1/RIP-1. p = 0.0003. Mann-Whitney test. **(I)** Negative stain electron microscopy of RAD-51 filaments formed after 5 min at 25 °C in the presence or absence of 250 nM (in nucleotides) 150-mer ssDNA. Quantification represents normalised fold increase of number of RAD-51 filaments relative to ‘RAD-51+ssDNA only’ condition. Six fields of view across different sections of the EM grid were evaluated and plotted. p = 0.0022. Mann-Whitney test.

Unexpectedly, direct visualisation of RFS-1/RIP-1(A647) molecules revealed that they bind to RAD-51^f^ clusters with extremely short dwell-times (Fig. 4E, F). Colocalization of RFS-1/RIP-1(A647) with RAD-51^f^ clusters (Fig. 4G) is consistent with recognition of nascent RAD-51 filaments rather than ssDNA. Collectively, our data indicate that RFS-1/RIP-1 promotes assembly of RAD-51 filaments by shutting down disassembly events at filament ends without being a stable component of DNA-bound RAD-51^f^ clusters. Hence, RFS-1/RIP-1 acts in a transient manner as a dynamic RAD-51 filament ‘chaperone’.

Rapid photobleaching and step-finding analysis (Autour et al., 2018; Watkins and Yang, 2005) allowed us to calibrate the imaging system, determine the number of fluorophores in individual nucleating clusters and assess RAD-51 cluster growth (Fig. S6A, B). In accordance with a power-law fit, most RAD-51^f^ clusters are dimeric (Fig. S6C). Inclusion of RFS-1/RIP-1(A647) shifts the cluster population towards smaller species with the monomer fraction corresponding to ∼50% of the molecules. These data remain consistent with our power-law fit, k_obs_=J[RAD-51]^n^, where n = 1.6, indicating monomers, in addition to dimers, represent the substantial fraction of minimal nucleating units. Consistent with RFS-1/RIP-1(A647) stabilising RAD-51^f^ on ssDNA, we observe more frequent growth in the number of RAD-51^f^ protomers in individual clusters (Fig. 4H). The increased frequency of RAD-51^f^ cluster growth provides an explanation for the stimulation of filament growth rates observed with RFS-1/RIP-1.

To further validate that these growing RAD-51^f^ clusters indeed correspond to nascent RAD-51 filaments, we performed negative-stain electron microscopy (EM) using 150-mer ssDNA. Under the same buffer and protein-concentration used in single-molecule assays, EM analysis revealed that the inclusion of RFS-1/RIP-1 leads to enhanced RAD-51 filament formation when compared to RAD-51 alone (Fig. 4I). We propose that transient engagement of RFS-1/RIP-1 with RAD-51 filaments shuts down RAD-51 dissociation events at filament ends and shifts protomer addition-dissociation equilibrium towards higher net filament growth rates.

### ATP hydrolysis regulates transient engagement of RFS-1/RIP-1 with 5’ filament ends

We hypothesized that the short dwell-times of RFS-1/RIP-1 might result from ATP hydrolysis by RFS-1 or RAD-51 given that ATP hydrolysis releases Rad51 from DNA (Chi et al., 2006; Gataulin et al., 2018; Robertson et al., 2009). Indeed, upon inclusion of the slowly-hydrolysable ATP-analogue, ATP-γ-S, the dwell-time of RFS-1/RIP-1(A647) increased in ‘dipping’ experiments (Fig. S7A, B, C), indicating that ATP hydrolysis is at least partially responsible for RFS-1/RIP-1(A647) dissociation from RAD-51 filaments.

To further examine the dynamics properties of RFS-1/RIP-1, we first performed real-time imaging of RFS-1/RIP-1(A647) association with RPA-eGFP coated ssDNA in the presence of RAD-51 (Fig. 5A, B). Individual RFS-1/RIP-1(A647) molecules display very short dwell-times on ssDNA with ATP (median of 11.3 s, 9.5 - 15.8 s, 96% CI; Fig. 5A), similar to the dwell-times observed using the ‘dipping’ protocol described above. Upon inclusion of ATP-γ-S, the dwell-time of RFS-1/RIP- 1(A647) increased (median of 28.3 s; 22 s - 56.2 s, 97.6% CI; Fig. 5A). Notably, dwelling RFS-1/RIP-1(A647) molecules are frequently (71% of events scored, n = 26) located at the border of RPA-eGFP (blue) and RAD-51 filaments (dark; Fig. 5B). This corroborates our previous negative stain EM data (Taylor et al., 2015) that RFS-1/RIP-1 engages with 5’ filament ends (Taylor et al., 2016), but not within RAD-51 filaments.

**Figure 5.**
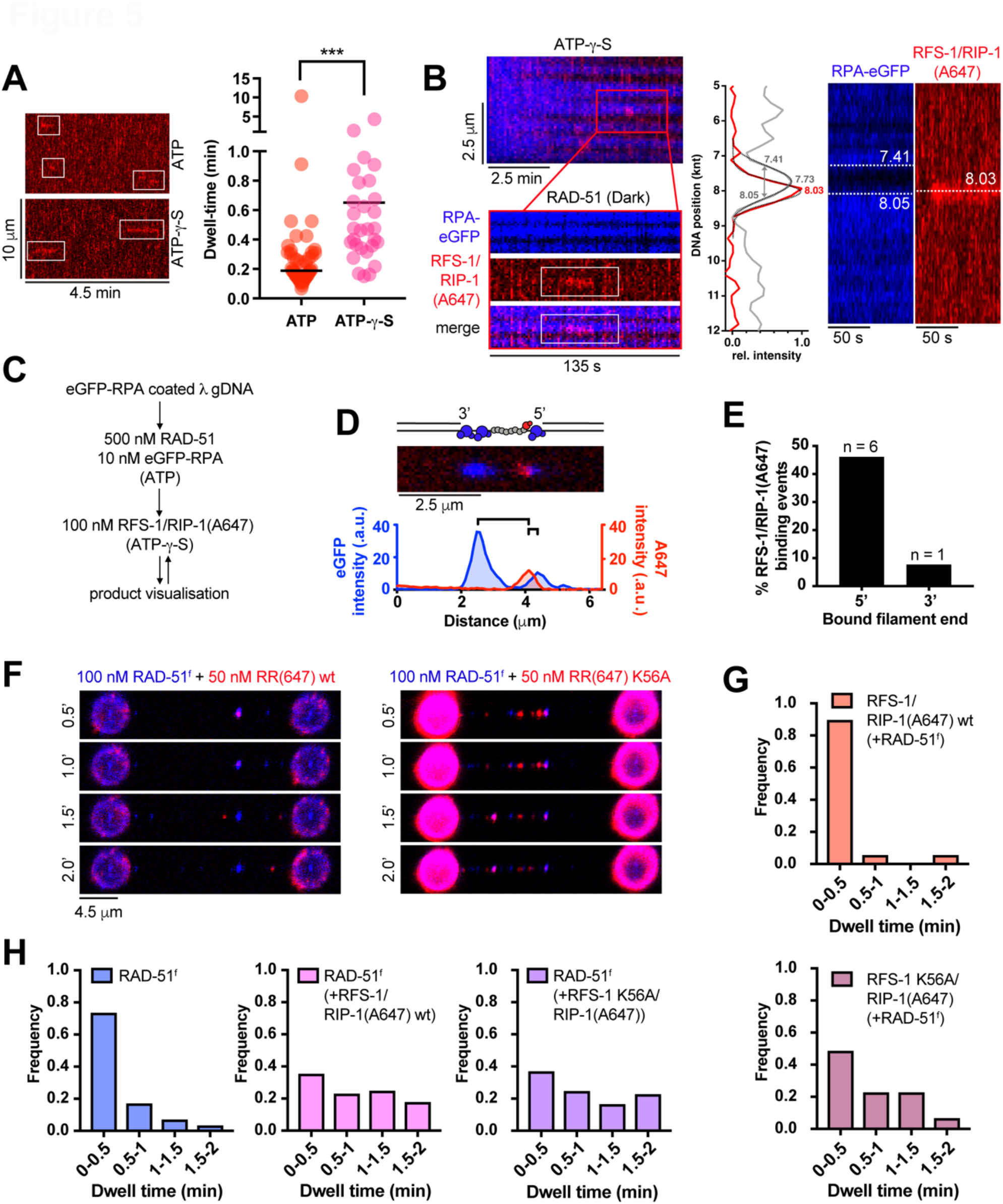
Transient engagement of RFS-1/RIP-1 with 5’ filament ends is mediated by ATPase activity. **(A)** Representative kymographs of dwelling single RFS-1/RIP-1(A647) (2.5 nM) complexes on RPA-eGFP coated ssDNA in the presence of 500 nM RAD-51 and ATP or ATP-γ-S (left panel). Quantification of experiment shown in left panel, for ATP n = 47 and ATP-γ-S n = 29. Black line represent median. p < 0.0001. Mann-Whitney test. (right panel) **(B)** Representative traces of single RFS-1/RIP-1(A647) complexes binding to RAD-51 filaments in ATP-γ-S. RPA-eGFP shown in blue, RAD-51 dark, RFS-1/RIP-1(A647) red. Quantification of the RFS-1/RIP-1(A647) end binding using custom position analysis algorithm. 2D scan showing RFS-1/RIP-1(A647) (red channel) binding to the RAD-51 filament (dark, blue channel) in the presence of RPA-eGFP (blue channel). To obtain the exact location of the RAD-51 paralog with respect to the RAD-51 filament, the centre of the filament is resolved first by fitting the reversed eGFP intensity (grey peak). Grey arrow, representing the peak’s width, marks the filament’s edges. Gaussian fit of the RFS-1/RIP-1(A647) intensity (red peak, red channel) indicates that the paralog binds to the periphery of the RAD-51 filament. **(C)** Protocol designed to visualise RFS-1/RIP-1(A647) binding to RAD-51 filaments grown on gapped λ DNA. **(D)** 2D scan of representative RAD-51-DNA complex obtained using protocol described in Fig. 5C. Proximity of A647 signal maximum to eGFP signal maximum on either 3’ or 5’ RAD-51 filament border was used to estimate 3’ or 5’ filament end binding polarity. **(E)** Quantification of filament 5’ or 3’ end binding frequencies by RFS-1/RIP-1(A647) (n=13). Remainder of RFS-1/RIP-1(A647) binding events observed in the experiment was binding to ‘dark’ RAD-51 filaments forming on extended dsDNA portion of the gapped molecule. No binding of RFS-1/RIP-1(A647) to RAD-51-free RPA-eGFP-coated ssDNA was observed. **(F)** Representative fluorescence images taken after RAD-51^f^+RFS-1/RIP-1(A647) or RAD-51^f^+RFS-1 K56A/RIP-1(A647) incubation–detection cycles with the same ssDNA construct; cumulative incubation time is indicated. RAD-51^f^ signal shown in blue. RFS-1/RIP-1(A647) signal shown in red. **(G)** Histograms of dwell times of wt RFS-1/RIP-1(A647) (n = 19; top) or RFS-1 K56A/RIP-1(A647) (n = 31; bottom) in the presence of RAD-51^f^. Number of dwell-time categories was adjusted to accommodate lower stability of bare ssDNA in the presence of RFS-1 K56A/RIP-1(A647). **(H)** Histograms of dwell times of RAD-51^f^ alone (n = 160; left) or in the presence of RFS-1/RIP-1(A647) wt (n = 57; middle), or dwell times of RAD-51^f^ in the presence of RFS-1 K56A/RIP-1(A647) (n = 49; right).

We established that RAD-51 filaments grow preferentially in 3’ to 5’ direction in the presence of RFS-1/RIP-1. We have also shown that RFS-1/RIP-1 shuts down dissociation of RAD-51 clusters on ssDNA leading to increase of their dwell-times and growth frequencies. This prompted us to investigate whether labelled RFS-1/RIP-1 directly binds to 5’ filament ends using gapped DNA substrates with defined polarity. To directly visualise RFS-1/RIP-1 at filament ends, we incubated RAD-51 filaments with 100 nM RFS-1/RIP-1(A647) (Fig. 5C), which revealed proximity to 5’, but not 3’, RAD-51 filament ends (Fig. 5D, E). Collectively, these data confirm RFS-1/RIP-1 transiently binds 5’ filament ends and shuts down RAD-51 dissociation to allow for efficient filament growth in a 3’ to 5’ direction.

To determine whether the putative ATPase activity of RFS-1/RIP-1 is critical for the dissociation from RAD-51 filament ends, we performed ‘dipping’ experiments using Alexa 647-labelled RFS-1 K56A/RIP-1 complex, in which the lysine residue of the Walker A box of RFS-1 is mutated to alanine. Mutating lysin to alanine/arginine in recombinase Walker A motif was previously shown to lead to abolishment of ATP binding/hydrolysis *in vitro* (Chi et al., 2006). Strikingly, RFS-1 K56A/RIP-1(A647) displays dramatically increased dwell-times on RAD-51 ssDNA clusters when compared to wt RFS-1/RIP-1(A647) (Fig. 5F, G). Notably, RFS-1 K56A/RIP-1(A647) retains the ability to stabilise RAD-51^f^ clusters on ssDNA (Fig. 5H), which we attribute to its ability to bind RAD-51 clusters. Inclusion of RFS-1 K56A/RIP-1 also stimulates RAD-51 assembly on RPA-eGFP-coated ssDNA, albeit less efficiently than wt RFS-1/RIP-1 (Fig. S8A, B). When compared to wt RFS-1/RIP-1, RAD-51 filaments display slower growth rates in the presence of RFS-1 K56A/RIP-1 (Fig. S8C). These observations indicate that the RFS-1 K56A/RIP-1 mutant is effective at filament end-binding and stabilization of nucleating RAD-51 clusters, but is compromised for its ability to stimulate RAD-51 filament growth. Notably, increasing wt RFS-1/RIP-1 concentrations to near stoichiometric levels with RAD-51 compromises filament assembly and hinders RAD-51 filament growth when compared to the growth stimulation observed with sub-stoichiometric levels of wt RFS-1/RIP-1 (Fig. S8A, B, C). This suggests that excessive filament end binding at high concentrations of RFS-1/RIP-1 or Walker A box mutants in RFS-1 blocks further recruitment of RAD-51 protomers and hinders filament elongation. We also verified filament end binding by RFS-1 K56A/RIP-1 on filaments formed by Cy3-labelled RAD-51 (RAD-51^Cy3^; Fig. S8D, E). The increased dwell-times of the RFS-1 K56A/RIP-1 mutant on RAD-51 filaments are also in agreement with yeast two hybrid analysis where a robust interaction is detectable between RFS-1 K56A/RIP-1 mutant and RAD-51, but not between wt RFS-1/RIP-1 and RAD-51 (Fig. S8F). Furthermore, RFS-1 K56A/RIP-1 and RFS-1 K56R/RIP-1 both form a super-shifted complex with RAD-51-ssDNA in bulk EMSA assays, whereas wt RFS-1/RIP-1 does not (Fig. S8G). Collectively, these results indicate that wt RFS-1/RIP-1 engages transiently with RAD-51 filaments, whereas the higher residence time of RFS-1 K56A/RIP-1 on RAD-51 filament ends results in the formation of aberrant co-complexes, which hinder filament growth (Fig. S8C).

### RFS-1 K56A/R variants are competent for RAD-51 stabilization, but inefficient for HR *in vivo*

To determine whether the dynamic engagement of RFS-1/RIP-1 during RAD-51 filament growth plays an important role in HR *in vivo*, we generated nematode knock-in mutant strains for both RFS-1 K56A and RFS-1 K56R using the CRISPR-Cas9 system (Fig. 6A). Similar to the *rfs-1* (null) deletion strain, *rfs-1* K56A and *rfs-1* K56R mutant strains display a modest increase in the levels of embryonic lethality compared to the N2 (wt) strain without significant brood size reduction (Fig. 6B, C). In agreement with previous reports (Taylor et al., 2015; Ward et al., 2007), the *rfs-1* null strain displays sensitivity to agents that induce replication fork breakage (Ward et al., 2007). Similarly, *rfs-1* K56A and K56R Walker A mutants are sensitive to cisplatin (CDDP), camptothecin (CPT) and nitrogen mustard (HN2), although to a lesser extent than *rfs-1* null animals. Sensitivity to hydroxyurea (HU) is not observed in the *rfs-1* K56A and *rfs-1* K56R mutant strains (Fig. 6D).

**Figure 6.**
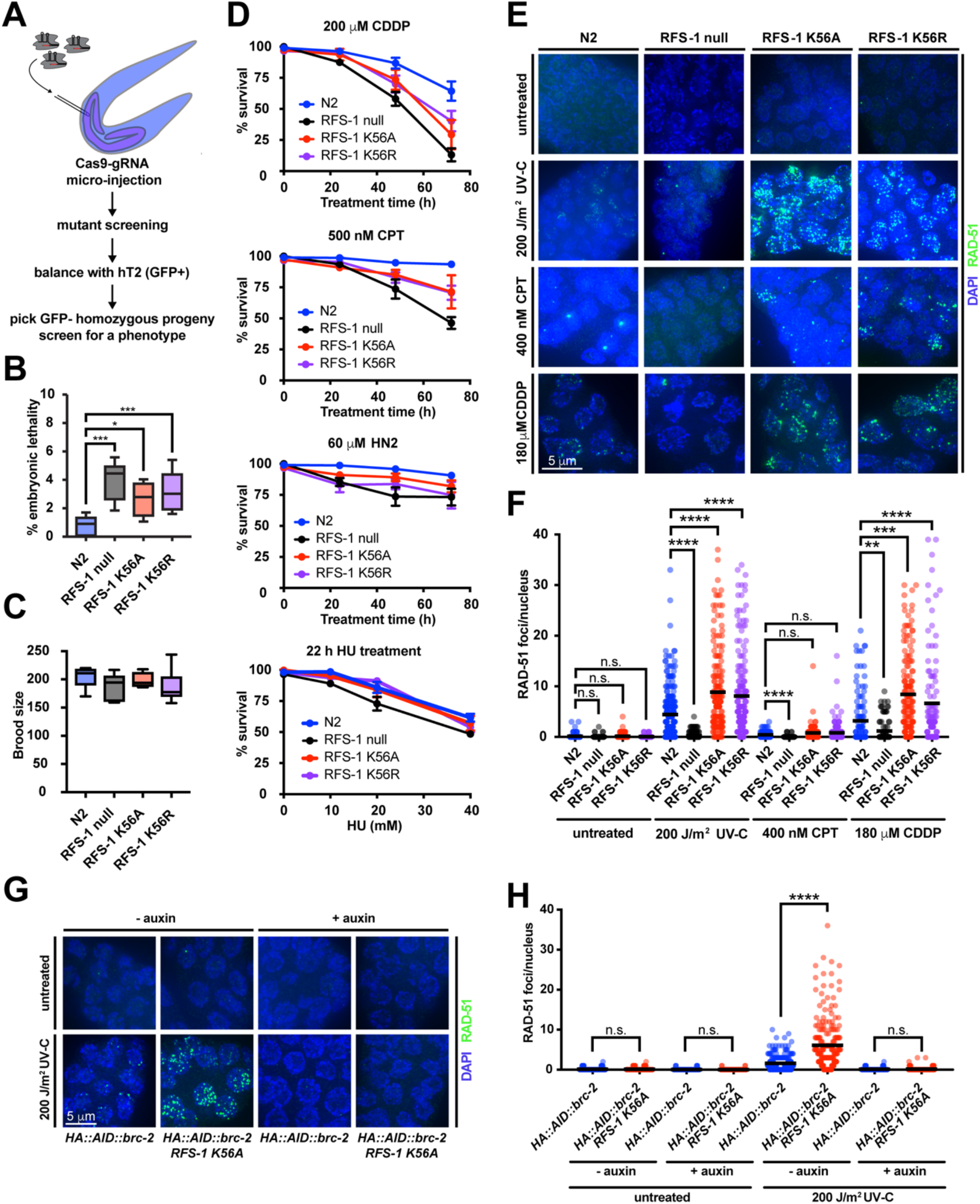
RFS-1 K56A/R *C. elegans* strains accumulate HR-incompetent RAD-51 foci after DNA damage. **(A)** A schematic for CRISPR-Cas9 knock-in strategy for generation of RFS-1 Walker box mutant strains. **(B)** Embryonic lethality analysis of *rfs-1* mutants *in vivo*. Percentage of unhatched eggs was scored after 24 hours in strains of the indicated genotype. Progeny of 6-8 worms were evaluated. **(C)** Brood size of strains of the indicated genotype. Progeny of 5-8 worms were evaluated. **(D)** The indicated strains were treated with indicated doses of genotoxins for the indicated time. *rfs-1* null, *rfs-1 K56A* and *rfs-1 K56R* strains display increased sensitivity to replication–associated DSBs lesions caused by CDDP, CPT and HN2 mustards, while sensitivity to replication fork stalling in the presence of hydroxyurea is mild in *rfs-1* null strain and absent in RFS-1 K56A and RFS-1 K56R strains. Error bars represent SEM. **(E)** Representative images of the mitotic compartment of *C. elegans* germline after treatment with different genotoxins. DAPI staining (blue), RAD–51 staining (green). Scale bar represents 5 μm. **(F)** Quantification of RAD-51 focus formation in the mitotic zone of the worm germline under different treatments in strains of the indicated genotype. Between 99 and 261 cells were quantified for each condition in 2-3 representative germlines for each genotype. Mann-Whitney test was used for statistical analysis. **(G)** Representative images of the mitotic compartment of *C. elegans* germline of indicated genotypes after treatment with indicated dose of UVC grown in the presence or absence of auxin. HA::AID::brc-2 corresponds to *C. elegans* strain modified to express BRC-2 N-terminally fused to HA followed by auxin-inducible degron. DAPI staining (blue), RAD–51 staining (green). Scale bar represents 5 μm. **(H)** Quantification of RAD-51 focus formation in the mitotic zone of the worm germline under different treatments in strains of the indicated genotype. Between 99 and 261 cells were quantified for each condition in 3 representative germlines for each genotype.

Mitotic zones of extruded germlines were then examined for RAD-51 focus formation before and after CDDP, UV-C and CPT treatment. In agreement with previous studies (Ward et al., 2007), RAD-51 forms damage-induced foci in N2(wt) strains but not in the *rfs-1* null strain. Strikingly, *rfs-1* K56A and K56R Walker A mutant strains display extensive accumulation of RAD-51 foci in response to DNA damage (Fig. 6E, F), which persist into meiosis I. These data, together with DNA damage sensitivity, indicate that RAD-51 nucleates and is stabilized on DNA in the *rfs-1* K56A and *rfs-1* K56R mutant strains, but the resulting RAD-51 species are non-functional for repair of DNA damage via HR.

Since the RFS-1 mutants inefficiently disengage from filament ends *in vitro*, we considered the possibility that failure to efficiently disassembly from 5’ filament ends may result in short, but stable filaments unable to efficiently perform DNA strand exchange. Under these conditions, BRC-2 may continue loading RAD-51 on over-resected DNA (Symington, 2016) leading to the observed phenotype. Consistent with this possibility, the extensive accumulation of RAD-51 complexes in treated germlines of *rfs-1* K56A mutants is abolished by depletion of BRC-2 (Fig. 6G, H). Hence, BRC-2 promotes RAD-51 nucleation on RPA-coated ssDNA and RFS-1/RIP-1 acts downstream to stabilize and facilitate growth of nascent RAD-51 filaments *in vivo*, as observed in our single-molecule experiments.

In summary, our single-molecule and genetic data support a model where BRC-2 facilitates RAD-51 nucleation on RPA-coated ssDNA, with the recruitment of RAD-51 protomers in equilibrium with disassembly bursts from nascent RAD-51 filament ends. To shift the equilibrium in favour of filament growth, RFS/RIP-1 function sequentially as a molecular ‘chaperone’ by transiently binding RAD-51 filaments and preventing RAD-51 dissociation from 5’ filament ends, leading to stabilisation of RAD-51 on RPA-coated ssDNA. Subsequent release of RFS-1/RIP-1, which depends on ATP hydrolysis and the Walker box in RFS-1, allows for further addition of RAD-51 protomers at 5’ filament end leading to the formation of a RAD-51 ssDNA filament proficient for strand exchange (Fig. 7).

**Figure 7.**
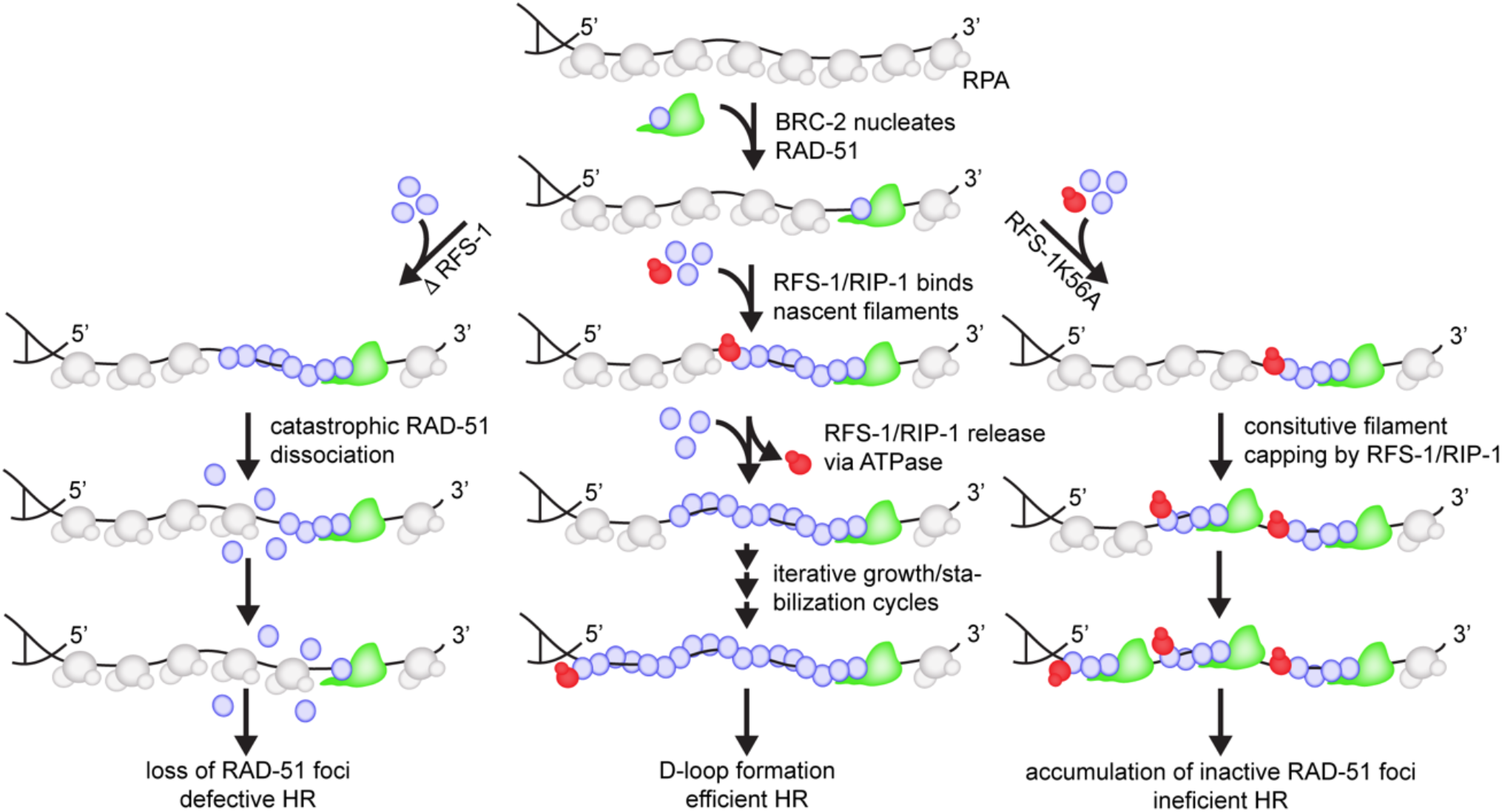
Model describing the mechanism of metazoan RAD-51 presynaptic filament assembly. BRC-2 nucleates RAD-51 on RPA-coated resected ssDNA. Nascent RAD-51 filaments are bound at 5’ end by RFS-1/RIP-1 which stabilizes growing RAD-51 filaments preventing burst-like disassembly from filament ends. ATPase mediated dissociation of RFS-1/RIP-1 allows for further recruitment of RAD-51 protomers allowing filament growth in 5’ direction. Inability of RFS-1 K56A/RIP-1 to dissociate from bound RAD-51 filaments results in extensive end-capping and formation of very stable shorter less active RAD-51 filaments, which accumulate by the action of BRC-2-mediated RAD-51 nucleation.

## DISCUSSION

In bacteria, RecA filaments form on ssDNA via a two-step mechanism: rate-limiting nucleation and rapid elongation (Bell et al., 2012). The RecFOR mediator complex enhances both steps (Bell et al., 2012), but whether a similar mechanism applies to eukaryotic mediators is not known. Here, we establish the mechanism of eukaryotic Rad51 filament nucleation and growth, which requires the sequential action of mediator proteins. We implicate BRC-2 primarily as a nucleation factor that promotes RAD-51 accumulation on RPA-coated ssDNA. In contrast to RecA, eukaryotic Rad51 nuclei grow very slowly alone. However, inclusion of the Rad51 paralog complex, RFS-1/RIP-1, promotes rapid growth of RAD-51 filaments in a 3’→5’ direction. This filament growth stimulation requires highly dynamic 5’ filament end binding by RFS-1/RIP-1, which is regulated by intrinsic ATP hydrolysis. Hence, RFS-1/RIP-1 acts as a classical ‘chaperone’ in mediating the growth of functional RAD-51 filaments *in vitro* and *in vivo*.

### Division of labour between mediator proteins

Our single-molecule data reveal for the first time the contribution of mediator proteins to nucleation and growth of RAD-51 filaments. The division of labour between BRC-2 and RFS-1/RIP-1 likely stems from their intrinsic biochemical properties: BRC-2 possesses high affinity for both ssDNA and RAD-51 in solution (Martin et al., 2005; Petalcorin et al., 2006); while RFS-1/RIP-1 displays very low affinity for ssDNA and only interacts with RAD-51 when bound to ssDNA (Taylor et al., 2015). This is consistent with RFS-1/RIP-1 acting on RAD-51 filaments, and in agreement with multiple cellular studies showing Rad51 paralogs act downstream of BRCA2 (Chun et al., 2013; Jensen et al., 2013). Consistent with this notion, recruitment of yeast (Lisby et al., 2004) and vertebrate (Raschle et al., 2015) Rad51 paralogs (Rad55-Rad57) to DSBs is Rad51-dependent, while DNA damage-dependent nematode BRC-2 foci accumulate independently of RAD-51 (Martin et al., 2005). In line with cellular data, *in vitro* studies have shown that inclusion of full-length human BRCA2 increases the number of RAD51 filaments, but not their length (Shahid et al., 2014), whereas, addition of budding yeast Rad51 paralogs Rad55-Rad57 increases the length of Rad51 filaments in negative stain EM (Liu et al., 2011). Hence, our findings, supported by previous observations, establish that BRCA2 acts first to promote nucleation, followed by Rad51 paralogs acting on ssDNA-bound Rad51 to promote filament growth.

Interestingly, the division of labour between BRC-2 and RAD-51 paralogs is not absolute, as we found that BRC-2 also significantly stimulates RAD-51 filament growth, albeit to a much lower extent than RFS-1/RIP-1. It remains to be tested if interactions of BRCA2 with other HR factors (e.g. PALB2 or DSS1) might further stimulate filament growth enhancement by BRCA2, particularly given that both PALB2 and DSS1 enhance BRCA2’s stimulatory effect in bulk DNA strand exchange assays (Buisson et al., 2010; Zhao et al., 2015). Rad51 paralogs also appear to enhance RAD-51 nucleation rates, although this is likely due to stabilization of small RAD-51 clusters on ssDNA rather than through direct nucleation.

### Directionality of RAD-51 filament growth

Bacterial RecA filaments were previously shown to grow bi-directionally with a two-fold kinetic preference for the 5’→3’ direction (Bell et al., 2012). Using asymmetrically positioned gapped DNA substrates, we have shown that, contrary to RecA, nematode RAD-51 filaments grow very slowly in both directions. Strikingly, addition of RFS-1/RIP-1 specifically increases growth rates in a 3’→5’ direction, opposite to RecA. Given that human BRCA2 recognizes the 3’ filament interface (Pellegrini et al., 2002) and full-length human BRCA2 binds at the 3’ filament end (Shahid et al., 2014), we propose that following RAD-51 nucleation by BRC-2, the free 5’-end is engaged and stabilized by RFS-1/RIP-1, allowing nascent RAD-51 filaments to be extended in a 3’→5’ direction. Furthermore, engagement of opposing ends of the filament by recombination mediators also allows for efficient cooperation during filament assembly, potentially explaining the observed synergy between BRC-2 and RFS-1/RIP-1.

### Rad51 paralogs “chaperone” Rad51 filament growth

Previous work using bulk EM imaging postulated that Rad51 paralogs stabilize Rad51 filaments by stably intercalating into them (Liu et al., 2011). In contrast, direct single molecule imaging revealed that RFS-1/RIP-1 engages with 5’ RAD-51 filaments ends in a highly dynamic manner. The 5’ filament end binding is in line with predictions from previous bulk experiments (Taylor et al., 2016). Modulating RAD-51 through filament ends would explain the requirement for sub-stoichiometric RFS-1/RIP-1 concentrations for RAD-51 assembly rate stimulation in bulk assays (Taylor et al., 2015). Previous work has established that disassembly from Rad51 filament ends occurs by catastrophic dissociation bursts involving multiple Rad51 protomers (van Mameren et al., 2009). A similar phenomenon was observed in our ‘dipping’ experiments. On the other hand, recombinase filaments grow slowly where monomer addition is rate-limiting (Bell et al., 2012; Joo et al., 2006). Supressing costly dissociation bursts, as observed in the ‘dipping’ experiments, would make an effective regulatory mechanism to stimulate net RAD-51 filament growth or stabilize short unstable RAD-51 clusters.

We further establish that this rapid turnover is dependent on ATPase activity of RFS-1 and demonstrate that RFS-1 Walker box mutations or blocking ATP hydrolysis stabilizes RAD-51 on ssDNA, but failure to disengage RFS-1/RIP-1 from 5’ filament ends hinders further filament growth. Taken together, our results suggest that RFS-1/RIP-1 acts as RAD-51 filament ‘chaperone’ by recognizing nascent 5’ RAD-51 filament ends, transiently binding them to prevent filament disassembly, and dissociating to permit further Rad51 protomer addition and filament growth. Iterative cycles allow for efficient filament extension similarly to how classical molecular chaperones mediate unfolded protein recognition, folding, and release cycles.

This model is consistent with our phenotype analysis of *rfs-1* null and Walker A box K56A/R mutant strains. Loss of RFS-1 in the null strain confers marked sensitivity to DNA damaging agents accompanied by the loss of RAD-51 foci. Strikingly, in *rfs-1* Walker A box K56A/R knock-in strains, DNA damage sensitivity is also observed, but RAD-51 foci extensively accumulate, indicative of stabilized but HR-incompetent RAD-51 complexes. Extensive RAD-51 foci observed in *rfs-1* K56A/R background may result from continued resection and RAD-51 loading by BRC-2 as perturbations of downstream HR are known to increase resection tract length (Chung et al., 2010; Haas et al., 2018; Symington, 2016). Consistently, BRC-2 depletion abolishes the accumulation of RAD-51 foci in *rfs-1* K56A/R mutant strain, demonstrating that BRC-2 functions prior to RFS-1/RIP-1 *in vivo*.

Collectively, our findings disprove previous models that postulated that Rad51 paralogs intercalate and stably associate with Rad51 filaments (Liu et al., 2011) and thereby act as roadblocks for filament disruption by the anti-recombinase Srs2 helicase. Our model is corroborated in an accompanying paper by Roy et al., showing that yeast Rad55-Rad57 also exhibits highly dynamics engagement with DNA-bound Rad51 complexes during filament formation on ssDNA curtains and this depends on ATP hydrolysis. Further support to a ‘chaperone’ model of Rad51 paralog action is evident from direct imaging of Srs2 performed in the same study, which revealed eviction of residual Rad55-Rad57 bypassing Srs2 molecules without any reduction in velocity directly, thereby disproving the roadblock model. On the contrary, Rad55-Rad57 stimulates Rad51 filament formation behind translocating Srs2 molecules in the presence of free Rad51.

The notion that Rad51 paralogs are not a stable components of Rad51 complexes is further supported by work in Chinese hamster ovary cells, where different Rad51 paralogs were shown to turnover rapidly from the sites of replication upon challenge with genotoxic agents, while Rad51 persists at these sites significantly longer (Somyajit et al., 2015). Walker A box mutants of different Rad51 paralogs significantly decreased the turnover of these proteins. A similar behaviour has been proposed for full-length human BRCA2 *in vitro* (Shahid et al., 2014). Together with our findings, these observations imply that mediator proteins may act as transient Rad51 filament binding factors to promote efficient Rad51 filament assembly without stable association with it.

### HR and replication fork protection *in vivo*

Loss of Rad51 (Hashimoto et al., 2010), BRCA2 (Schlacher et al., 2011) and, more recently, Rad51 paralogs (Somyajit et al., 2015) have been shown to induce Mre11-dependent degradation of replication tracts upon replication fork stalling induced by HU and accumulation of ssDNA gaps at and behind replication forks (Hashimoto et al., 2010). Human RAD51 K133R forms very stable (Chi et al., 2006), albeit HR-incompetent filaments (Stark et al., 2004). Expression of this mutant can suppress fork degradation phenotypes observed in BRCA2 deficient cells, suggesting a genetic separation from HR (Schlacher et al., 2011). Notably, RFS-1 and RIP-1 deficient nematode strains display strong phenotypes in response to replication fork-blocking lesions, but not in IR-induced DNA breaks or meiotic recombination (Ward et al., 2007). Interestingly, while RFS-1 loss sensitizes nematodes to CDDP, CPT and HU, *rfs-1* K56A/R strains are sensitive to CDDP and CPT, while in response to HU, almost a full rescue is observed. This might imply that similarly to Rad51 K133R, RAD-51 filaments constitutively bound and stabilized by RFS-1 K56A/R are still capable of supporting fork protection upon HU treatment, but are defective for conventional HR repair of CPT and CDDP-induced DSBs.

In conclusion, the work presented here, together with the accompanying study by Roy et al., reveals a highly conserved mechanism of Rad51 filament assembly through the action of filament ‘chaperones’.

## STAR ⋆ METHODS

- KEY RESOURCES TABLE
- LEAD CONTACT AND MATERIALS AVAILABILITY
- EXPERIMENTAL MODEL AND SUBJECT DETAILS
- METHOD DETAILS
  • Protein expression, purification and yeast two-hybrid interaction assays. Purification of Rad51, RPA, Srs2, Hed1 and Dmc1
  • Fluorescent labelling of RAD-51 and RFS-1/RIP-1. IR sensitivity assays
  • Single molecule imaging using dual optical trapping system
  • Single-step photobleaching and image analysis
  • EMSA
  • D-loop formation assay
  • Oligonucleotide-based DNA strand exchange assay
  • Negative stain electron microscopy
  • Genome editing using CRISPR-Cas9 in C. elegans
  • Injection mix for CRISPR-Cas9 editing
  • Auxin-inducible protein degradation
  • Treatment of C. elegans with genotoxic agents
  • Immunostaining and image acquisition
- QUANTIFICATION AND STATISTICAL ANALYSIS
- DATA AND CODE AVAIBILITY

## LEAD CONTACT AND MATERIALS AVAILABILITY

Further information and requests for resources and reagents should be directed to and will be fulfilled by the lead contact, Simon J Boulton.

## METHOD DETAILS

### Protein expression, purification and yeast two-hybrid interaction assays

RAD-51, RFS-1/RIP-1 complex and RPA complex were expressed and purified as described previously (Taylor et al., 2015; Taylor and Yeeles, 2018). Human eGFP-RPA was a kind gift from Mauro Modesti (CRCM, Marseille). Yeast two-hybrid was performed as described previously (Taylor et al., 2015). Codon-optimized BRC-2 ORF was cloned into pET MBP-1a His_6_-MBP-BRC-2 (referred to as BRC-2 in the manuscript) was expressed in BL21(DE3) *E. coli* strain at 17 °C overnight using 0.1 mM IPTG for the induction of protein expression. Cells were lysed in Lysis Buffer (25 mM Tris-HCl pH 7.5, 500 mM KCl, 10% glycerol, 1 mM DTT, 0.01% NP40 substitute, cOmplete EDTA-free protease inhibitor tablets (1/50 ml), cat no. 11873580001, Roche). After sonication and centrifugation at 20,000 rpm for 1h, clarified lysate was applied to Ni-NTA (nitrilotriacetic acid, Qiagen) resin for 1.5h, washed with Lysis buffer and Lysis buffer containing 20 mM Imidazole. Proteins were eluted using Elution buffer 500 (25 mM Tris-HCl pH 7.5, 300 mM KCl, 10% glycerol, 0.5 mM EDTA, 1 mM DTT, 0.01% NP40 substitute, 200 mM Imidazole). Sample was then directly applied to amylose resin and allowed to be bound for 1h, amylose beads were washed with Wash buffer 300 (25 mM Tris-HCl pH 7.5, 300 mM KCl, 10% glycerol, 0.5 mM EDTA, 1 mM DTT, 0.01% NP40 substitute). Protein was eluted using Elution buffer 300 (25 mM Tris-HCl pH 7.5, 300 mM KCl, 10% glycerol, 0.5 mM EDTA, 1 mM DTT, 0.01% NP40 substitute, 30 mM maltose) and diluted two times with Elution buffer lacking KCl and maltose). Sample was then loaded onto pre-equilibrated HiTrap SP column, column was washed with 10 column volumes of Buffer A (25 mM Tris-HCl pH 7.5, 150 mM KCl, 10% glycerol, 0.5mM EDTA, 1 mM DTT, 0.01% NP40 substitute) and eluted using linear salt gradient (0-80%) of Buffer B (25 mM Tris-HCl pH 7.5, 1000 mM KCl, 10% glycerol, 0.5 mM EDTA, 1 mM DTT, 0.01% NP40 substitute). Fractions containing BRC-2 were pooled, concentrated, frozen and subsequently checked for purity using SDS-PAGE.

### Fluorescent labelling of RAD-51 and RFS-1/RIP-1

RAD-51 was labelled using amine-reactive FAM, Cy5 and Cy3 NHS-esters as described previously for RecA with modifications (Amitani et al., 2010). Briefly, protein storage buffer was exchanged using Zeba Column (0.5 mL resin, 3KDa MWCO) for labelling buffer (50 m*M* K_2_HPO_4_/KH_2_PO_4_ (pH 7.0), 200 mM KCl, 0.1 mM DTT, and 25% glycerol). Dyes were diluted in dry DMSO to 50 mM. Dyes and protein were mixed to final concentration of 50 µM protein and 500 µM FAM-SE or 150 µM Cy3/Cy5-NHS. Incubation on rotary shaker at 4 °C followed for 2 h 45 min (FAM-SE) or 2h (Cy5, Cy3-NHS). Reaction was terminated by the addition of Tris–HCl (pH 7.5) to a final concentration of 50 mM. Proteins were then buffer exchanged at least twice into storage buffer (50 mM Tris-HCl, pH 7.5, 300 mM KCl, 1 mM DTT, 0.01% NP40 substitute, 0.1 mM EDTA, 10% glycerol). Protein concentration was estimated by Coomasie staining and dye concentration was measured spectrophotometrically. Presence of minimum free dye concentration was assessed using SDS-PAGE on labelled proteins. Protein to dye concentration ratio was consistently 0.8-1.0.

For RFS-1/RIP-1 labelling we genetically fused ybbr tag (DSLEFIASKLA) on the C-terminus of RIP-1 downstream of 3xFLAG tag, separated by GGGSGGG linker. Proteins were expressed and purified using a previously established protocol (Yin et al., 2006). The labelling followed a protocol described elsewhere (Lim et al., 2017). Plasmid for Sfp expression was a kind gift from Dr. Meindert Lamers (LUMC). Sfp transferase was expressed and purified as described previously (Yin et al., 2006). The purified protein complexes (5 µM) were then labelled with CoA-Cy3 (30 µM) using recombinant Sfp phosphopantetheinyl transferase (1 µM) in final buffer condition of 50 mM HEPES pH 7.5, 300 mM KCl, 10 mM MgCl_2_, 1 mM DTT. After overnight incubation at 4 °C, the labelled protein complex was purified away from Sfp and free dye using Zeba column gel filtration system (0.5 mL resin, 50.000 MWCO). Proteins were stored in 20 mM Tris-acetate (pH 8.0), 100 mM potassium acetate, 10% glycerol, 1 mM EDTA, 0.5 mM DTT and subjected to SDS-PAGE and the fluorescent gel was scanned with a Typhoon9500 Scanner.

### Single molecule imaging using dual optical trapping system

Experiments were performed using commercially available C-trap (LUMICKS) setup. Protein channels of the microfluidics chip were first passivated with BSA (0.1% w/v in PBS) and Pluronics F128 (0.5% w/v in PBS), minimum 500 μl of both flowed through prior to use. Biotinylated ssDNA precursor was prepared as described previously (Candelli et al., 2014). To generate gapped λ DNA, biotinylated hairpin oligonucleotides were annealed to λ dsDNA ends and ligated (King et al., 2019). S. p. Cas9 D10A nickase (IDT) bound to previously described (Sternberg et al., 2014) guide RNAs were subsequently used to generate targeted DNA nicks. The reaction was then stored at 4 °C and directly diluted in PBS on the day of the experiment. DNA was captured between 4.5 -μm SPHERO Streptavidin Coated polystyrene beads at 0.005% w/v using the laminar flow cell, stretched and held at forces of 100 pN and higher until the strands were fully melted. The presence of ssDNA was verified by comparison to built-in freely joined chain model. For all the imaging conditions, ssDNA was held at forces between 10 and 20 pN, which corresponds roughly to 1.5-fold extension of B-form lambda dsDNA. Proteins were flown into incubation channels and bound to ssDNA by a previously-described dipping protocol. Importantly, under low-coverage regime (concentrations of 10-100 nM), a constant flow was kept during the incubations to minimize concentration variations due to surface adhesion of labelled proteins. Beads and DNA were kept in PBS during the experiment, while DNA was melted in 0.5xNTM buffer (25 mM Tris-HCl pH 7.5, 50 mM NaCl, 0.5 mM MgCl_2_) supplemented with 1mM ATP, oxygen scavenging system (2.5 mM 3,4-dihydroxybenzoic acid, 250 nM protocatechuate dioxygenase) and 0.2 mg/ml BSA. Proteins were flowed into the system in 1xNTM buffer (50 mM Tris-HCl pH 7.5, 100 mM NaCl, 1 mM MgCl_2_) supplemented with 1 mM ATP, oxygen scavenging system (2.5 mM 3,4-dihydroxybenzoic acid, 250 nM protocatechuate dioxygenase) and 0.2 mg/ml BSA. When high protein concentrations were used (≥500 nM), ATP-regeneration system consisting of 20 mM phospho-creatine and 20 μg/mL creatine kinase was also added into the reaction.

For ‘dipping assays’ performed with different cofactors, controls using AMP-PNP-Mg^2+^ and ADP-aluminium fluoride-Mg^2+^ were performed in addition to ATP-γ-S. However, no RFS-1/RIP-1(A647) binding to RAD-51^f^ clusters was observed under these conditions. BRC-2 was also assessed on bare ssDNA in the ‘dipping assay’, however, we observed formation of extremely bright aggregated GFP-BRC-2-ssDNA clusters on ssDNA containing numerous (>20) molecules of BRC-2, which was not the case under physiological conditions with RPA. For confocal imaging, three excitation wavelengths were used, 488 nm for eGFP and 6-FAM, 532 nm for Cy3 and 638 nm for Cy5, with emission detected in three channels with blue filter 512/25 nm, green filter 585/75 nm and red filter 640 LP. Imaging conditions for ‘dipping assay’: 15% laser power, 0.1ms/pixel dwell-time, 100 nm pixel size. Imaging conditions for ‘RPA-eGFP displacement assay’: 2% blue laser power, 5% red laser power, 0.1ms/pixel dwell-time, 100 nm pixel size, 1.5 s inter-frame wait time.

### Single-step photobleaching and image analysis

15% blue laser power was used to bleach RAD-51^f^ clusters in minimal imaging area to obtain sufficiently high bleaching time resolution. Scans were sectioned and stacked in Fiji using a custom-written script. Maximum likelihood estimation was used to determine each of the photobleaching steps within a maximum intensity/frame n. trace as previously described (Autour et al., 2018). The step sizes were subsequently binned and the histogram was fit to a double Gaussian equation in GraphPad Prism. 7. For cluster growth analysis, individual clusters were analysed for intensity increase in-between frames normalized to single-step intensity values. A cluster was considered as growing if the number of RAD-51^f^ promoters in the cluster increased by at least a single RAD-51^f^ protomer during the time the cluster dwelled on ssDNA. The growth frequency of RAD-51^f^ clusters was reported for each individual ssDNA molecule. For real-time RPA-eGFP displacement analysis, real-time force and fluorescence data were exported from Bluelake HDF5 files and analysed using custom-written scripts in Pylake Python package. Force was down sampled to 3 Hz for plotting. For RPA-eGFP free patch edge binding analysis, custom position-analysis script was built to extract the position of individual RPA-eGFP peaks and depressions, A647 intensity peaks were then aligned and their maxima position extracted to monitor proximity to the RPA-eGFP signal depression edges. Worm-like chain (WLC) model for λ dsDNA was used as a reference for force-extension curve comparison. Custom-written WLC fitting script was used to calculate contour length and subsequently gapped length of gapped DNA substrates. Growth rates in real-time experiments as well as dwell-times and binding frequencies were estimated in Fiji. Nucleation frequencies were plotted as a function of RAD-51^f^ concentration and fitted with power-law in GraphPad Prism 7. Dwell-times of RAD-51 clusters were binned into appropriate dwell-time categories cumulative survival analysis was performed using Igor 8.0. Mann-Whitney test was used to assess statistical significance of the data where appropriate.

### EMSA

Proteins were diluted from concentrated stocks into Storage Buffer, which was also used in no protein controls. For native polyacrylamide gels, proteins were mixed with a master mix (containing 60 nM (nucleotides) 5’-[32P]-labelled 60mer oligonucleotide (ACGCTGCCGAATTCTACCAGTGCCTTGCTAGGACATCTTTGCCCACCTGCAGGTTCACCC), 20 mM Tris-HCl (pH 7.5), 8% glycerol, 1 mM DTT, 50 mM sodium acetate, 2 mM MgCl_2_ and 2 mM ATP, and incubated for 10 min, before crosslinking with 0.25% glutaraldehyde for 10 min, all at 25°C. Reactions were resolved on 1% agarose gels in 1X TBE (70 V, 2 h 20 min). Gels were dried and imaged by autoradiography or using a storage phosphor screen (Amersham Biosciences) and Typhoon9500 and quantified using Fiji. For fluorescence experiments using RAD-51^f^ and/or labelled RFS-1/RIP-1 complex, proteins were incubated with 20 nM 49mer oligonucleotide AGCTACCATGCCTGCACGAATTAAGCAATTCGTAATCATGGTCATAGCT in 35 mM Tris-HCl (pH 7.5), 50 mM KCl, 1 mM DTT, 2 mM MgCl_2_, 2 mM ATP and incubated for 10 min at 25°C followed by resolution on 0.8% agarose gel in 1X TAE (70 V, 60 min). Gels were dried ad imaged uing Typhoon9500 and appropriate filter settings. % of DNA binding was assessed using Fiji.

### D-loop formation assay

RAD-51 and RAD-51^f^ were diluted from concentrated stocks into T Buffer (25 mM Tris-HCl (pH 7.5), 10% glycerol, 0.5 mM EDTA (pH 8.0), 100 mM KCl), which were also used in no protein controls. Proteins were mixed with 30 nM FAM-labelled 90mer ssDNA in 35 mM Tris-HCl (pH 7.5), 50 mM KCl, 1 mM DTT, 2 mM MgCl_2_ and 2 mM ATP and incubated for 10 min at 25 °C followed by addition of 0.54 μg pBS(-) dsDNA plasmid for further 15 min incubation at 25 °C. Reactions were terminated by SDS-PK treatment for 10 min at 37 °C. 90V/35 minute resolution using 1xTAE, 0.8% agarose gel electrophoresis followed. Gels were scanned using Typhoon9500 with appropriate filter settings.

### Oligonucleotide-based DNA strand exchange assay

40mer dsDNA was prepared by annealing 5’-fluorescein-labelled 40mer oligonucleotide (TAATACAAAATAAGTAAATGAATAAACAGAGAAAATAAAG) to the complementary unlabelled 40mer oligonucleotide (CTTTATTTTCTCTGTTTATTCATTTACTTATTTTGTATTA) in 50 mM Tris-HCl (pH 7.5), 100 mM NaCl, 10 mM MgCl_2_, and stored at stock concentration 200 nM (moles). Proteins were diluted from concentrated stocks into T Buffer (25 mM Tris-HCl (pH 7.5), 10% glycerol, 0.5 mM EDTA (pH 8.0), 50 mM KCl), which was also used in no protein controls. Proteins were mixed with 5.6 nM (moles) 150mer oligonucleotide (TCTTATTTATGTCTCTTTTATTTCATTTCCTATATTTATTCCTATTATGTTTTATTCATTTACTTATTCTTTATGTTCATTTTTTATATCCTTTACTTTATTTTCTCTGTTTATTCATTTACTTATTTTGTATTATCCTTATCTTATTTA), 50 mM Tris-HCl (pH 7.5), 1 mM DTT, 100 μg/ml of BSA, 2 mM ATP, 4 mM CaCl_2_ in 12.5 μl reaction volume at 25 °C for 10 min. 0.5 μl dsDNA stock and 0.5 μl 0.1 M spermidine were then added incubated for 1:30 h. The samples were deproteinized with 0.1% SDS and 12.5 μg proteinase K at 37 °C and resolved in 10% polyacrylamide gels in 1X TBE (80 V, 1 h 15 min). Gels were imaged on a Typhoon9500 and quantified using Fiji.

### Negative stain electron microscopy

RAD-51 and RFS-1/RIP-1 in indicated concentrations were incubated with 250 nM (in nucleotides) 150mer poly(dT) ssDNA in 50 mM Tris-HCl, pH 7.5, 100 mM NaCl, 2 mM MgCl_2_, 2 mM ATP for 5 min °C. For negative staining, Quantifoil R2/2, 2 nm carbon, 400 Cu mesh grids were glow discharged for 30 sec at 25 mA with a K100X glow discharger (EMS), 4 μl of sample was added to the grid left for 1 min. Excess sample was blotted away leaving a thin film. Then the grid was dipped into buffer solution twice and dipped twice into 2% uranyl acetate solution, blotting in between. Negative stain EM data were acquired on Tecnai™ Spirit TEM operated at 120 kV, equipped with a n FEI Eagle CCD camera.

### Genome editing using CRISPR-Cas9 in *C. elegans*

Genome editing by CRISPR-Cas9 was performed using preassembled Cas9-sgRNA complexes (trRNA, crRNA, Cas9) and single-stranded DNA oligos (used as repair templates) as described before(Paix et al., 2016). *dpy-10* was used as a co-injection marker to select progeny carrying Cas9-induced edits. The following sequences were used to generate crRNAs (IDT): *rfs-1* K56 mutants:

TTTAGGAGTTGGTAAAACAC; *HA::AID*::*brc-2*: TTTTTAGATGAGTCACCCAT; *dpy-10*: GCTACCATAGGCACCACGAG. The following repair templates were used, ordered as single-stranded DNA oligos at 4 nmol (IDT): *rfs-1K56A:* TTCATCCAGGAAAATGCTACGAAATTGATGGCGATCTGGGTGTAGGAGCTACGCAAGTATGAATTCATATATTTTATTTAGAGAATTTTCC;

*rfs-1K56R:* TTCATCCAGGAAAATGCTACGAAATTGATGGCGATCTGGGTGTAGGACGAACGCAAGTATGAATTCATATATTTTATTTAGAGAATTTTCC;

*HA::AID::brc-2* oligo 1: CAGACTTTACCAGAATATTGTGACATCGACCGATGTACCCATACGATGTTCCAGATTACGCTATGCCTAAAGATCCAGCCAAACCTCCGGCCAAGGCACAAGTTGTGGGATGGCCACCGGTGAGATCATACCGGAA;

*HA::AID::brc-2* oligo 2: GTTGTGGGATGGCCACCGGTGAGATCATACCGGAAGAACGTGATGGTTTCCTGCCAAAAATCAAGCGGTGGCCCGGAGGCGGCGGCGTTCGTGAAGGGTGACTCATCTAAAAAAGTGTTAGTCAAGATTTA;

*dpy-10:* CACTTGAACTTCAATACGGCAAGATGAGAATGACTGGAAACCGTACCGCATGCGGTGCC TATGG TAGCGGAGCTTCACATGGCTTCAGACCAACAGCCTAT.

### Injection mix for CRISPR-Cas9 editing

crRNAs and trRNA were reconstituted with nuclease-free duplex buffer to 200 µM and mixed in equal volumes to generate crRNA:trRNA duplex at 100 µM. Cas9/crRNA/trRNA complexes were generated by adding 2 µl of crRNA:trRNA duplex (100 µM) of the target gene, 0.2 µl of *dpy-10* crRNA:trRNA duplex (100 µM), and 2.95 µl of Cas9 nuclease V3 (at 61 µM, #1081059, IDT) and incubating the mix at room temperature for 5 minutes. The final injection mix was prepared by adding 0.6 µl of each ssDNA repair template from a 100 µM stock and 0.5 µl of *dpy-10* repair template (10 µM stock) to 5.15 µl of the Cas9/crRNA/trRNA complex, the mix was completed with H_2_O to obtain a final volume of 10 µl. The injection mix was directly injected into the gonads of young adult worms. Following injection, worms were placed onto individual NG agar plates seeded with *E. coli* (OP50) and incubated at 25 °C for three days. Roller and dumpy worms, caused by Cas9-dependent editing of the *dpy-10* gene, were picked individually to plates and allowed to produce progeny that was screened by PCR for the presence of the desired edit.

#### C. elegans strains generated in this study

|  |  |
| --- | --- |
| ATG563 | *rfs-1(fq127 [K56R]) III/hT2 (I;III)* |
| ATG567 | *rfs-1(fq130 [K56A]) III/hT2 (I;III)* |
| ATG600 | *brc-2(fq140[HA::AID::brc-2]) III; ieSi38 [P<sub>sun-1</sub>::TIR1::mRuby::sun-1 3' UTR, cb-unc-119(+)] IV* |
| ATG608 | *brc-2 (fq140 [HA::AID::brc-2]) rfs-1 (fq130 [K56A]) III; ; ieSi38 [P<sub>sun-1</sub>::TIR1::mRuby::sun-1 3' UTR, cb-unc-119(+)] IV* |

#### Other C. elegans strains used in this study

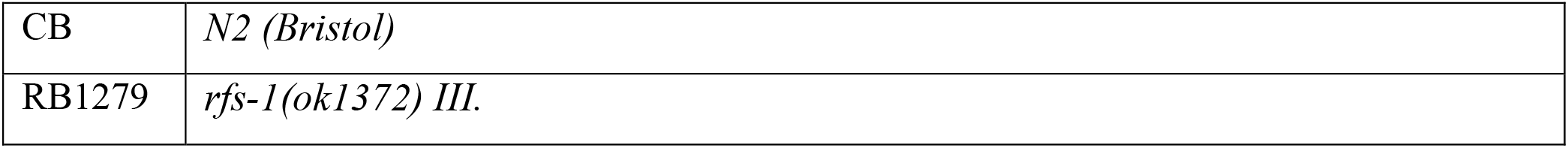

|  |  |
| --- | --- |
| CB | *N2 (Bristol)* |
| RB1279 | *rfs-1(ok1372) III.* |

### Auxin-inducible protein degradation

Auxin-mediated degradation of BRC-2 in the germline was performed by creating a strain homozygous for the *ieSi38* transgene, expressing *TIR-1-mRuby* under the *sun-1* promoter(Zhang et al., 2015), and for the CRISPR-generated *HA::AID::brc-2* allele. Young adult worms were placed on NG agar plates containing 4 mM auxin (Indole-3 acetic acid, Alfa Aesar, # A10556) seeded with *E. coli* (OP50) and allowed to lay eggs for two hours. Embryos were cultured on the auxin-containing plates for three days before young adults were picked and processed for immunostaining.

### Treatment of C. elegans with genotoxic agents

Exposure of worms to indicated doses of Cis-Diammineplatinum (II) dichloride (#P4394-250MG, Sigma, CDDP), Hydroxyurea (#H8627-5G, Sigma, HU), bis(2-chloroethyl)methylamine (122564-5G Sigma, HN2), and (S)-(+)-Camptothecin (#C9911-250MG, Sigma, CPT), was performed by placing worms on NG agar plates containing the desired amount of each genotoxic agent. Randomly picked young adult animals were placed on MYOB plates containing 200 μM CDDP, 500 nM CPT, 60 μM HN2 cisplatin or control plates. 3-5 worms were plated on each plate. Worms were moved every 24 hours to new drug-containing plates. Embryonic survival of progeny was then determined by determining the number of hatched eggs (calculated from initial number of layed eggs and dead eggs) on the 0-24, 24-48 and 48-72 hour plates. For HU treatment, worms were plated on plates containing indicated concentration of HU, for indicated period of time. Animals were transferred to HU–free plates and allowed to recover for 3 h. Worms were then allowed to lay eggs for 4 h. Dead eggs were counted 24 h after removing the parent animals.

### Immunostaining and image acquisition

Randomly picked gravid adult hermaphrodites were treated with cisplatin (CDDP, 180 μM) for 19 h and camptothecin (CPT, 500 nM) for 18 h in liquid culture Ionizing irradiation and UV-C (254 nm) treatment were performed on seeded plates. UV-C treatment of worms was performed on seeded plates using BLX-254 instrument. After treatment, animals were transferred to fresh seeded plates and allowed to recover (CDDP, 18 h; CTP, 7h; UV-C, 2h). Worms were washed twice in PBS, transferred to poly-L-lysine coated slides and germlines dissected. Germ lines from young adults hermaphrodites were dissected in egg buffer (118 mM NaCl, 48 mM KCl_2_, 2 mM CaCl_2_, 2 mM MgCl_2_, 5 mM HEPES at pH 7.4) and fixed in 1% paraformaldehyde containing 0.1% Tween for 5 min. Slides were frozen in liquid nitrogen, then immersed for 1 min in methanol at –20°C and transferred to PBST (1× PBS, 0.1% Tween). After washing the slides three times in PBST for 5 minutes, they were blocked in PBST 0.5% BSA for 30 minutes before incubating then overnight at room temperature with PBST containing anti-RAD-51 antibodies (a kind gift from A. Gartner) diluted 1:500 were incubated overnight at room temperature. Following three washes of 10 minutes each in PBST, slides were incubated with secondary antibodies (Alexa 488 α-rabbit, 1:500) for two hours in the dark. Following three washes of 10 minutes each in PBST, slides were counterstained with DAPI, washed in PBST for 1 hour and mounted using Vectashield. All images were acquired as stacks of optical sections with an interval of 0.2 μm using a Delta Vision deconvolution system equipped with an Olympus 1×70 microscope using 100xlens. Images were subjected to deconvolution using SoftWoRx 3.0 (Applied Precision).

## QUANTIFICATION AND STATISTICAL ANALYSIS

Scans were sectioned and stacked in Fiji using a custom-written script. Maximum likelihood estimation was used to determine each of the photobleaching steps within a maximum intensity/frame n. trace as previously described (Autour et al., 2018). The step sizes were subsequently binned and the histogram was fit to a double Gaussian equation in GraphPad Prism. 7. For cluster growth analysis, individual clusters were analysed for intensity increase in-between frames normalized to single-step intensity values. A cluster was considered as growing if the number of RAD-51^f^ promoters in the cluster increased by at least a single RAD-51^f^ protomer during the time the cluster dwelled on ssDNA. The growth frequency of RAD-51^f^ clusters was reported for each individual ssDNA molecule. For real-time RPA-eGFP displacement analysis, real-time force and fluorescence data were exported from Bluelake HDF5 files and analysed using custom-written scripts in Pylake Python package. Force was down sampled to 3 Hz for plotting. For RPA-eGFP free patch edge binding analysis, custom position-analysis script was built to extract the position of individual RPA-eGFP peaks and depressions, A647 intensity peaks were then aligned and their maxima position extracted to monitor proximity to the RPA-eGFP signal depression edges. Worm-like chain (WLC) model for λ dsDNA was used as a reference for force-extension curve comparison. Custom-written WLC fitting script was used to calculate contour length and subsequently gapped length of gapped DNA substrates. Growth rates in real-time experiments as well as dwell-times and binding frequencies were estimated in Fiji. Nucleation frequencies were plotted as a function of RAD-51^f^ concentration and fitted with power-law in GraphPad Prism 7. Dwell-times of RAD-51 clusters were binned into appropriate dwell-time categories cumulative survival analysis was performed using Igor 8.0. Mann-Whitney or other statistical test was used to assess statistical significance of the data where appropriate – sample size and statistical significance are indicated in the figures.

---

**Figure S1.**
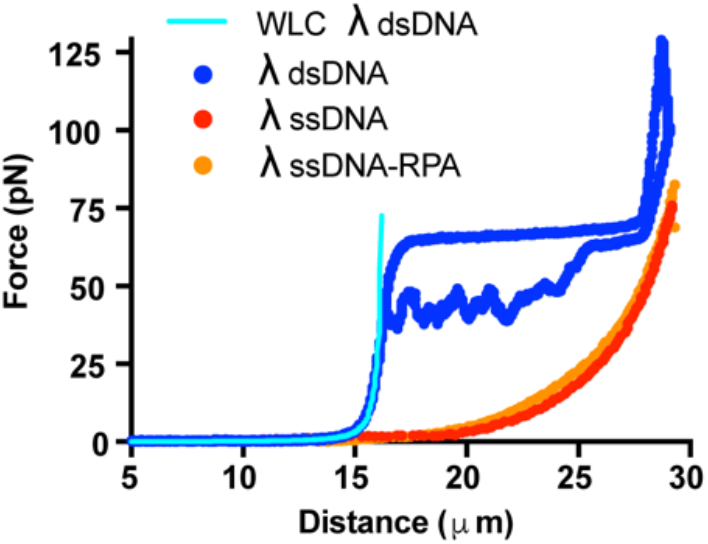
Force-extension curves recorded for different substrates. Force extension curves of 48.5 kb λ dsDNA (forward and reverse curve after partial melting), 48.5 kb λ ssDNA and 48.5 kb λ ssDNA coated by RPA. Blue line represents worm-like chain model (WLC) fit as a reference. RPA addition does not substantially change the shape of force-extension curve of λ ssDNA.

**Figure S2.**
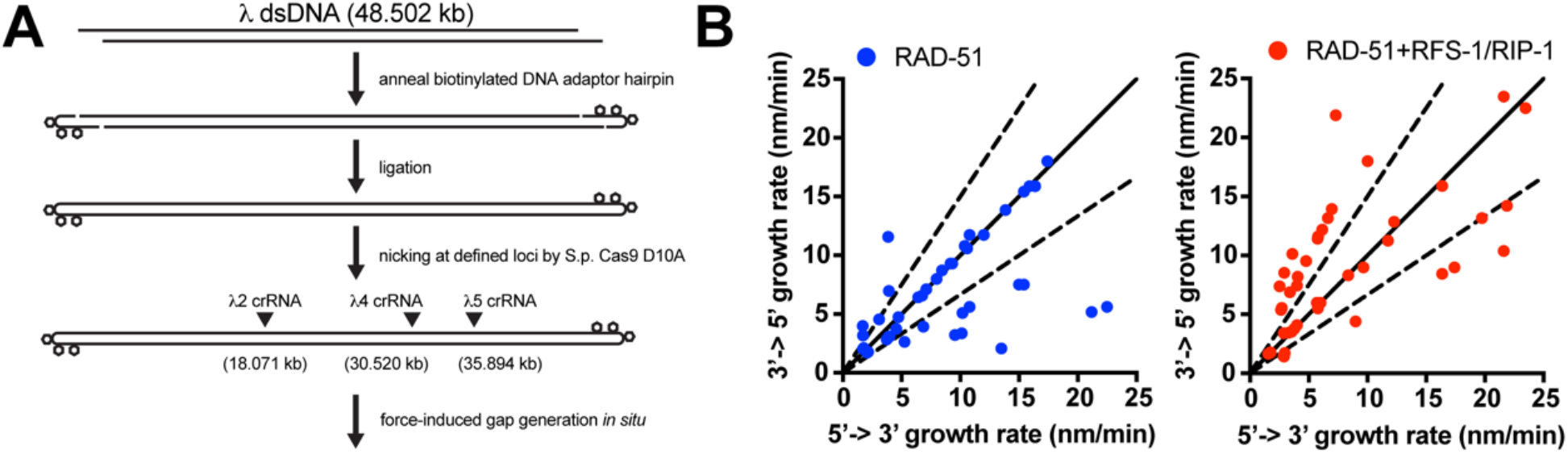
Generation of gapped DNA (gDNA) substrates and filament growth analysis. **(A)** A schematic of protocol designed to generate gapped λ DNA (gDNA) substrate. Two pairs of guide RNAs (λ4 crRNA and λ5 crRNA; λ2 crRNA and λ5 crRNA) and subsequent force-induced DNA melting *in situ* were utilized to generate 5.3 and 17.8 kb ssDNA gaps. **(B)** Scatterplot of 5’ to 3’ and 3’ to 5’ growth rates of individual RAD-51 clusters in the absence or presence of 10 nM RFS-1/RIP-1 using λ gDNA substrates. 10 nM free RPA-eGFP was included in a fraction of reactions performed and analysed to suppress excessive nucleation and obtain better-resolved growing clusters.

**Figure S3.**
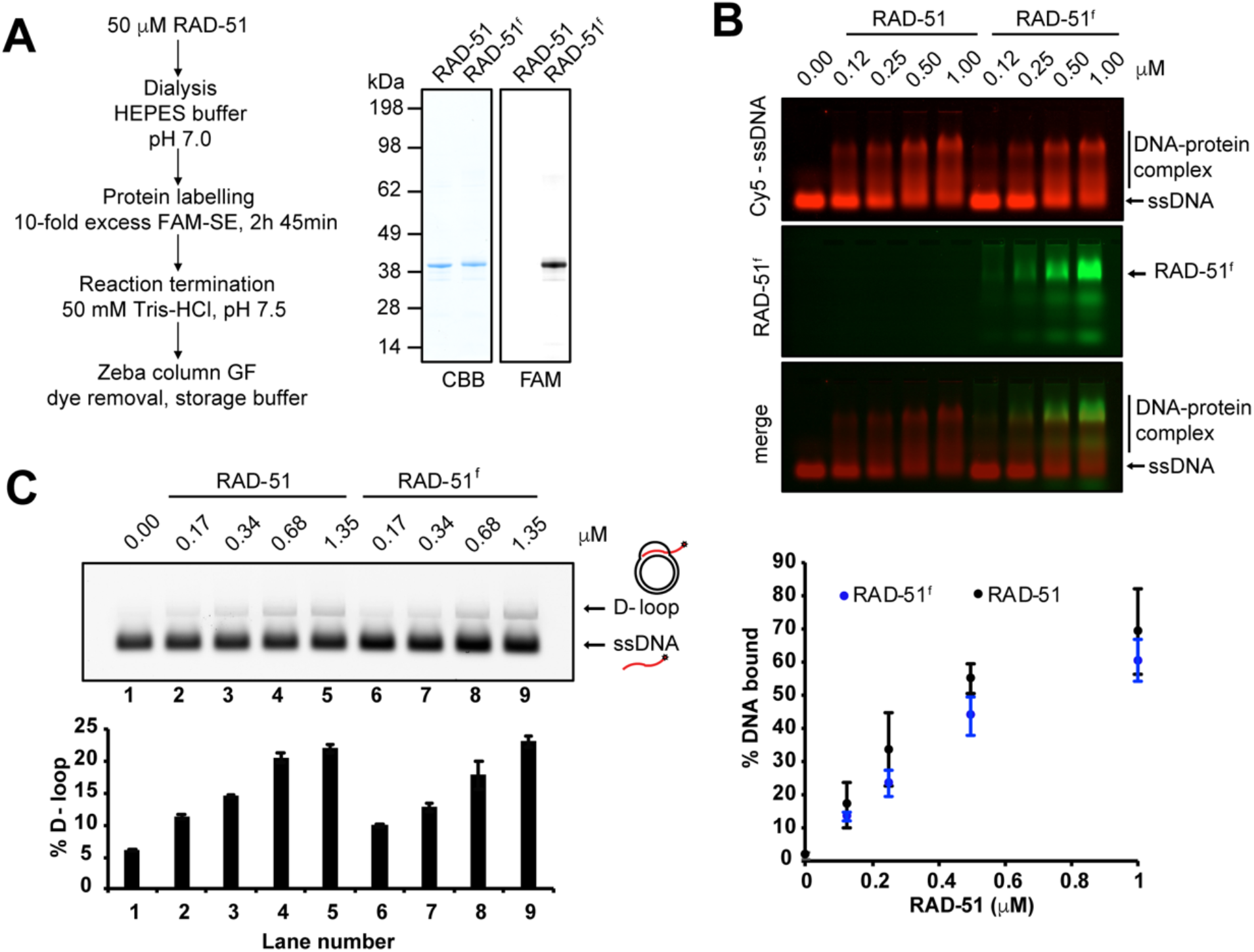
Fluorescent labelling of RAD-51. **(A)** Chemical labelling of RAD-51. RAD-51 was labelled in pH 7.0 using succinyl esters of 6-FAM (FAM-SE). After reaction termination and purification of labelled species, labelling efficiency was assessed and free dye component was evaluated using SDS-PAGE and subsequent fluorescent imaging. 1:1 labelling stoichiometry was achieved as measured spectrophotometrically. Higher labelling ratios attenuate RAD-51 DNA binding and D-loop formation activities. Proteins were labelled typically with 80-100% labelling efficiency. **(B)** EMSA showing that both RAD-51 and RAD-51^f^ bind ssDNA to a similar extent. Proteins were incubated with 20 nM (in molecules) FAM-labelled 49-mer ssDNA for 10 minute at 25 °C. DNA-protein complexes were resolved for 60 min at 70V in 0.8% agarose gel TAE electrophoresis at 4 °C. (n = 3). Error bars represent SD. **(C)** D-loop formation assay comparing DNA-pairing activities of RAD-51 and RAD-51^f^. Proteins were incubated with 30 nM (in molecules) FAM-labelled 90-mer ssDNA for 10 minute at 25 °C followed by addition of 0.54 μg pBS(-) plasmid DNA. Reactions were terminated by SDS-Proteinase K treatment for 10 min at 37 °C. DNA was resolved for 30 min at 90V in 0.8% agarose gel TAE electrophoresis at 4 °C. (n = 3). Error bars represent SD.

**Figure S4.**
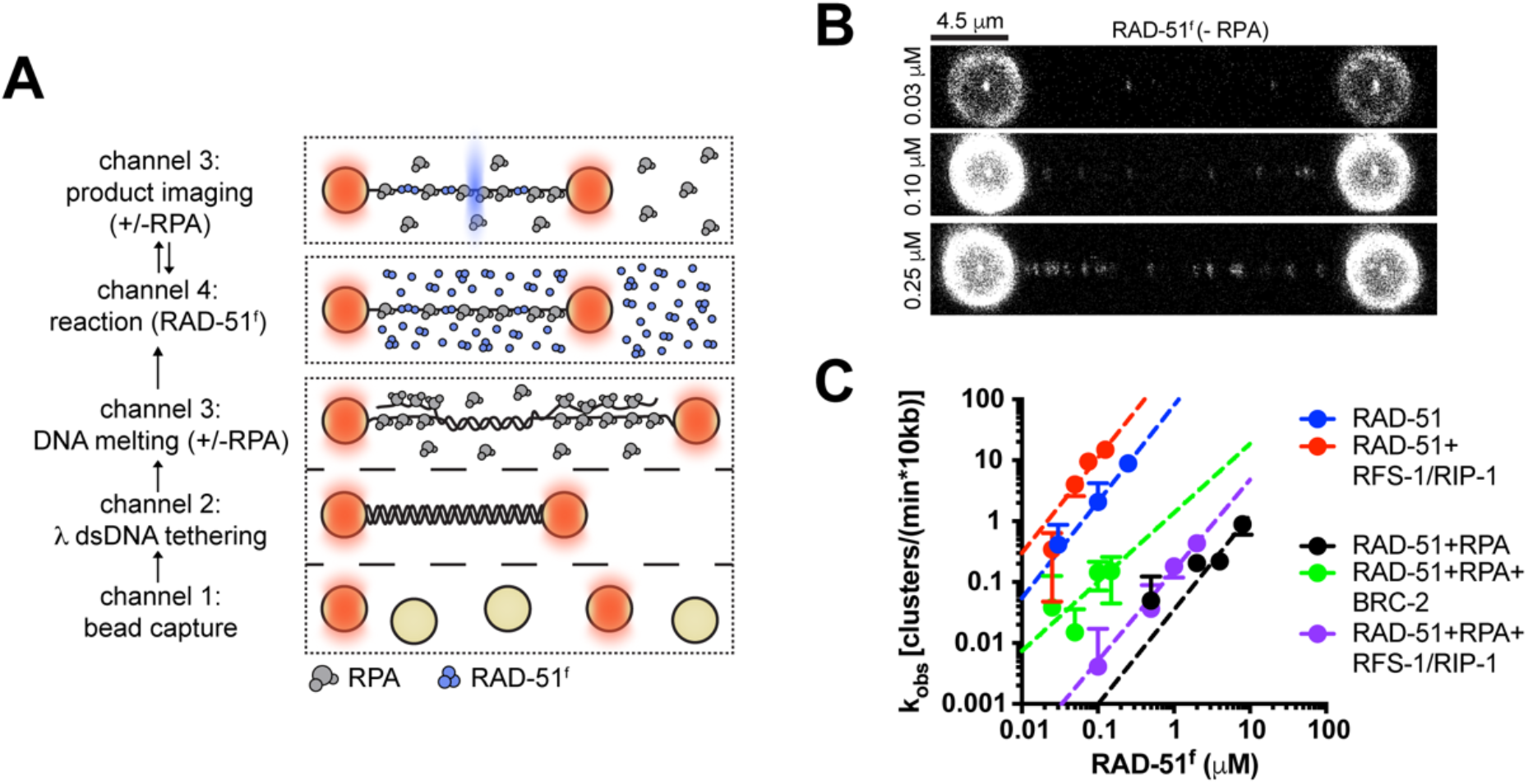
‘Dipping’ protocol to determine RAD-51 nucleation rates. **(A)** A four-channel microfluidics device was used for the assembly of the 5-step single-molecule binding experiments: (1) trapping of streptavidin-coated microspheres and (2) tethering of a single λ dsDNA between the spheres, (3) force-induced melting of dsDNA into ssDNA, (4) incubation in a flow channel with fluorescent RAD-51^f^ (blue) and ATP, and visualization of the RAD-51^f^ DNA complex in observation channel and buffer containing ATP. **(B)** Representative fluorescence images taken after 30s of RAD-51^f^ incubation– detection cycle with different RAD-51^f^ concentrations present in the solution. 4.5 µm scale bar. **(C)** RAD-51^f^ concentration dependence of nucleation rate. Dotted line represents power-law fit (*k*_*obs*_ = *J*[RAD-51]^*n*^) yielding an exponent *n* = 1.6 ± 0.2 for RAD-51^f^ in the absence of RPA (R^2^ = 0.72), *n* = 1.6 ± 0.2 for RAD-51^f^ in the absence of RPA and the presence of stoichiometric amounts of RFS-1/RIP-1 (R^2^ = 0.90), *n* = 1.6 ± 0.4 for RAD-51^f^ in the presence of RPA (R^2^ = 0.83), *n* = 1.1 ± 0.4 for RAD-51^f^ in the presence of RPA and stoichiometric amounts of BRC-2 (R^2^ = 0.21) and *n* =1.5 ± 0.1 for RAD-51^f^ in the presence of RPA and stoichiometric amounts of RFS-1/RIP-1 (R^2^ = 0.88). Error bars indicate SD.

**Figure S5.**
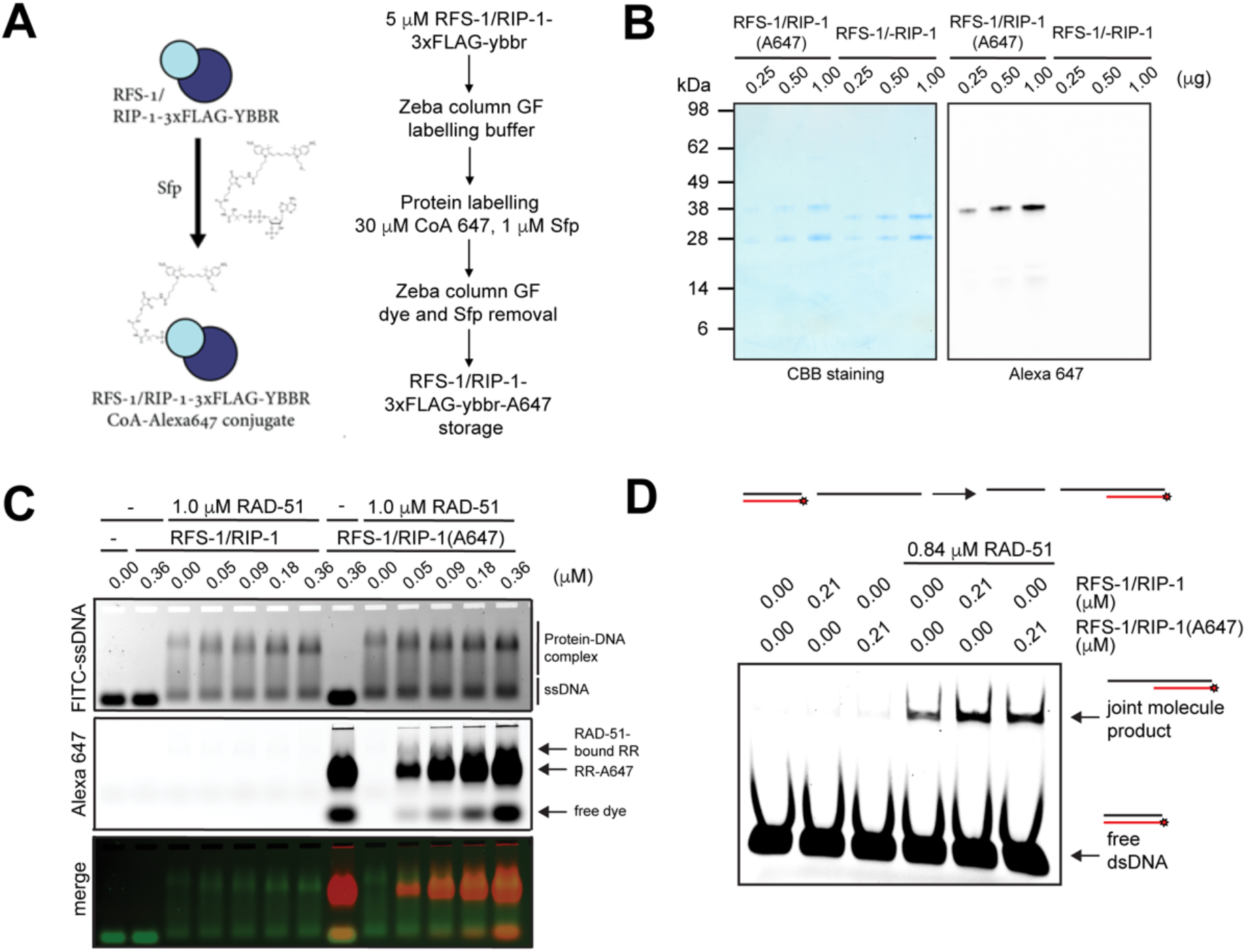
Fluorescent labelling of RFS-1/RIP-1. **(A)** Scheme and protocol for fluorescent labelling of ybbr-tagged RFS-1/RIP-1 complex. **(B)** Coomasie Brilliant Blue staining (left) and A647 fluorescence (right) of SDS-PAGE resolved RFS-1/RIP-1-3xFLAG or RFS-1/RIP-1-3xFLAG-ybbr-CoA-Alexa647 proteins. Loaded amount of protein is indicated. Proteins were labelled typically with 70-80% labelling efficiency. **(C)** EMSA showing that both RFS-1/RIP-1-3xFLAG and RFS-1/RIP-1-3xFLAG-ybbr-CoA 647 bind RAD-51-ssDNA complexes and induce a similar electrophoretic mobility shift. Proteins were incubated with 30 nM (in molecules) FAM-labelled 49-mer ssDNA for 10 min at 25 °C in the absence of glutaraldehyde crosslinking. DNA-protein complexes were resolved for 60 min at 70V in 0.8% agarose gel 1x TAE electrophoresis at 4 °C. **(D)** Strand exchange assay. Proteins were incubated for 10 min with 5.6 nM (in molecules) 150-mer oligonucleotide at 25 °C for 10 min. 8 nM (in molecules) dsDNA stock and 4 mM spermidine were then added followed by further incubation for 1.5 h at 25 °C. The samples were deproteinized with 0.1% SDS and 12.5 μg proteinase K at 37 °C and resolved using PAGE in 1x TBE.

**Figure S6.**
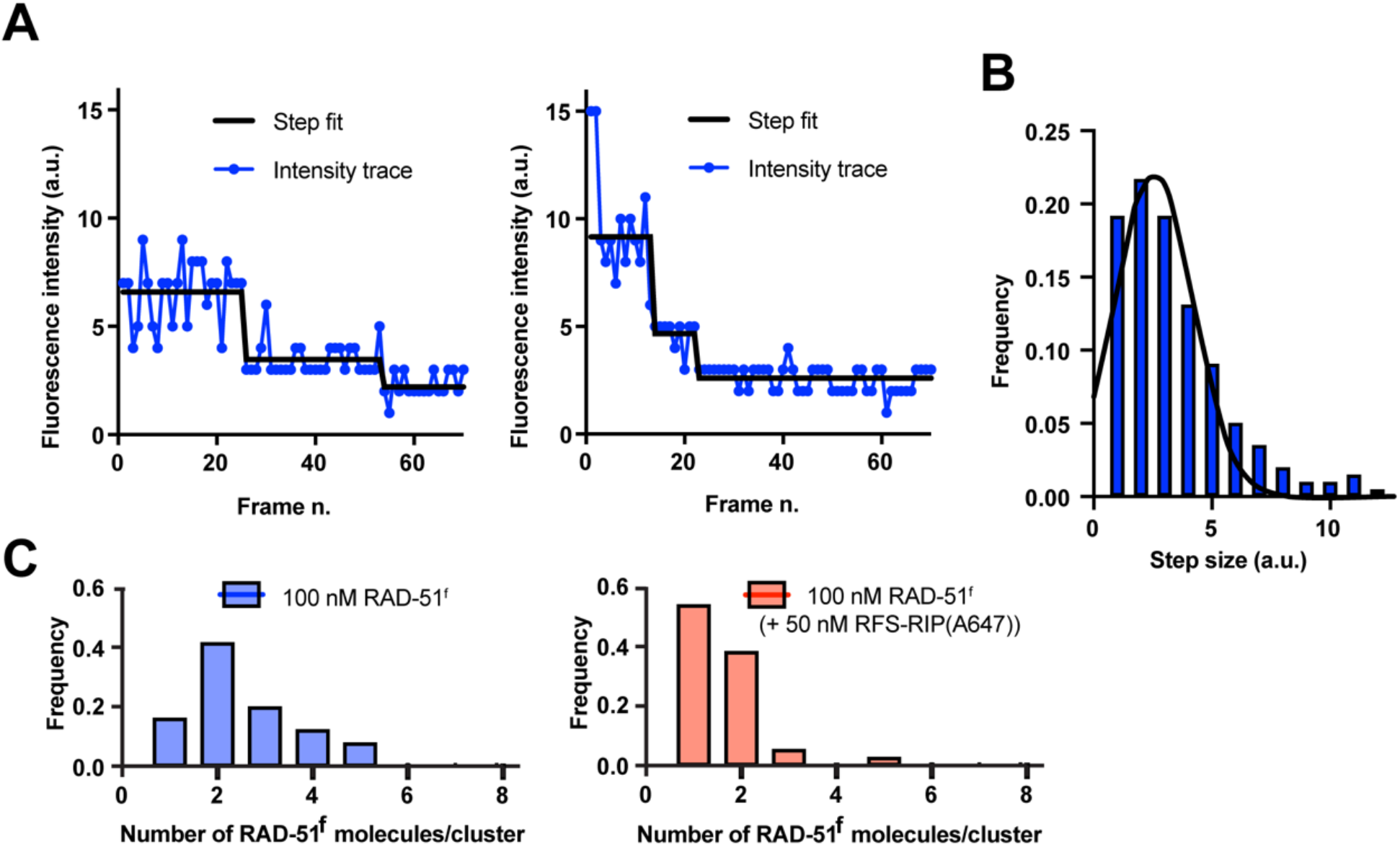
Single-step photobleaching analysis. **(A)** Examples of a fluorescence intensity trace of the RAD-51^f^ cluster during continuous photobleaching. Black line represents stepping fit of the trace. **(B)** Fluorescence intensity single-molecule calibration histogram for RAD-51^f^. The histogram shows the values of the fluorescence intensity trace of the single steps of multiple photobleaching traces (n = 198). Black line represents gaussian fit. Second two-step gaussian was excluded from the analysis. Mean = 2.55. S.D. = 1.67. R^2^ = 0.90. **(C)** Histogram of RAD-51^f^ cluster size in the presence (n = 76) or absence (n = 181) of RFS-1/RIP-1(A647).

**Figure S7.**
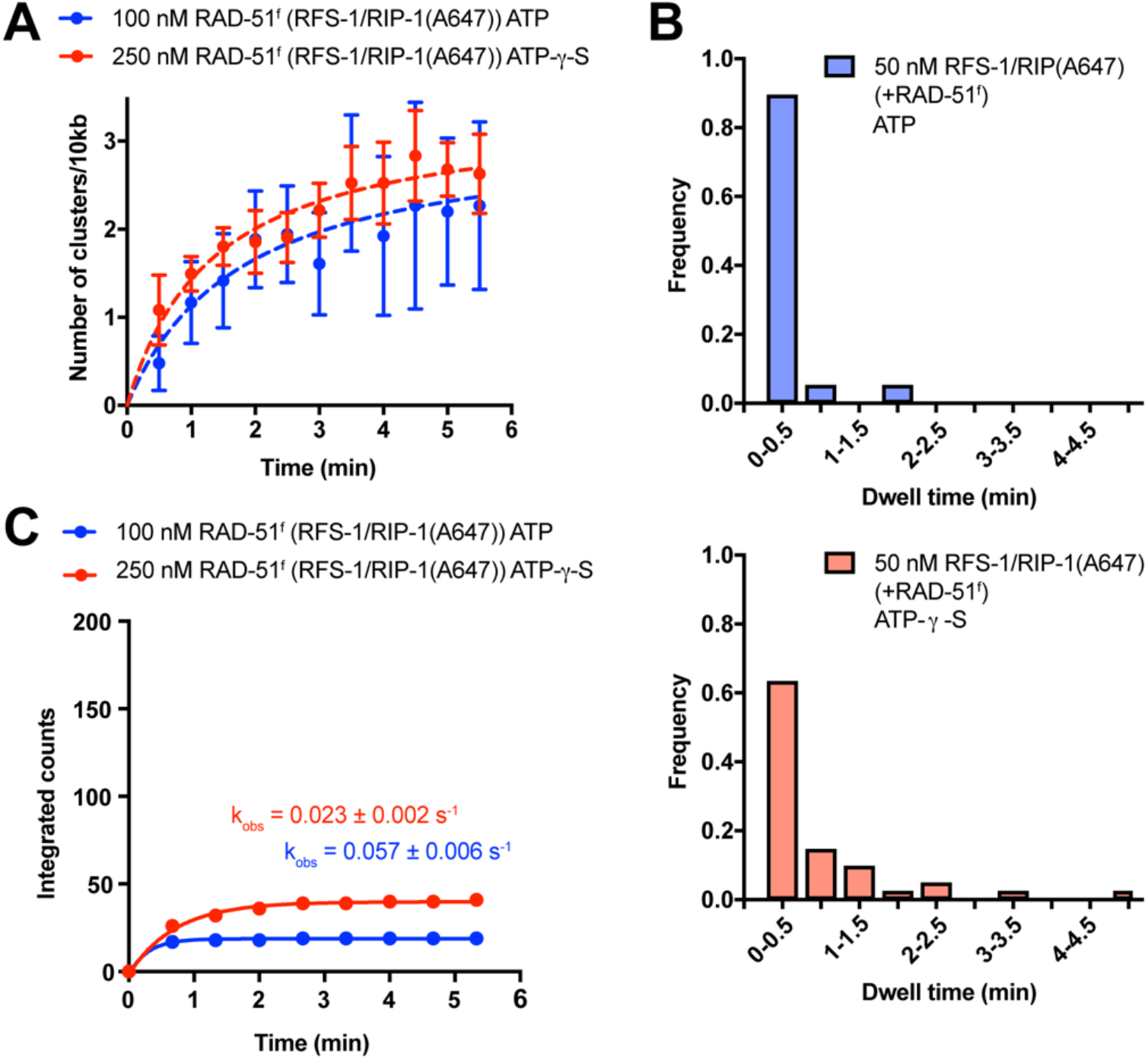
RFS-1/RIP-1(A647) dwell-times in the presence of ATP-γ-S. **(A)** Quantification of RAD-51^f^ nucleation frequency over time in the presence of ATP (n = 9) or ATP-γ-S (n = 9) and RFS-1/RIP-1(A647) in the absence of RPA. Exponential fits are displayed as dashed lines. Error bars represent SEM. RAD-51 concentrations were chosen to yield similar nucleation frequencies. In similar experiment, negligible nucleation frequencies were observed with 100 nM RAD-51 alone. **(B)** Histograms of dwell times of RFS-1/RIP-1(A647) in the presence of RAD-51^f^ with ATP or ATP-γ-S. **(C)** Cumulative survival plots of data presented in 6b. Lines represent exponential fits.

**Figure S8.**
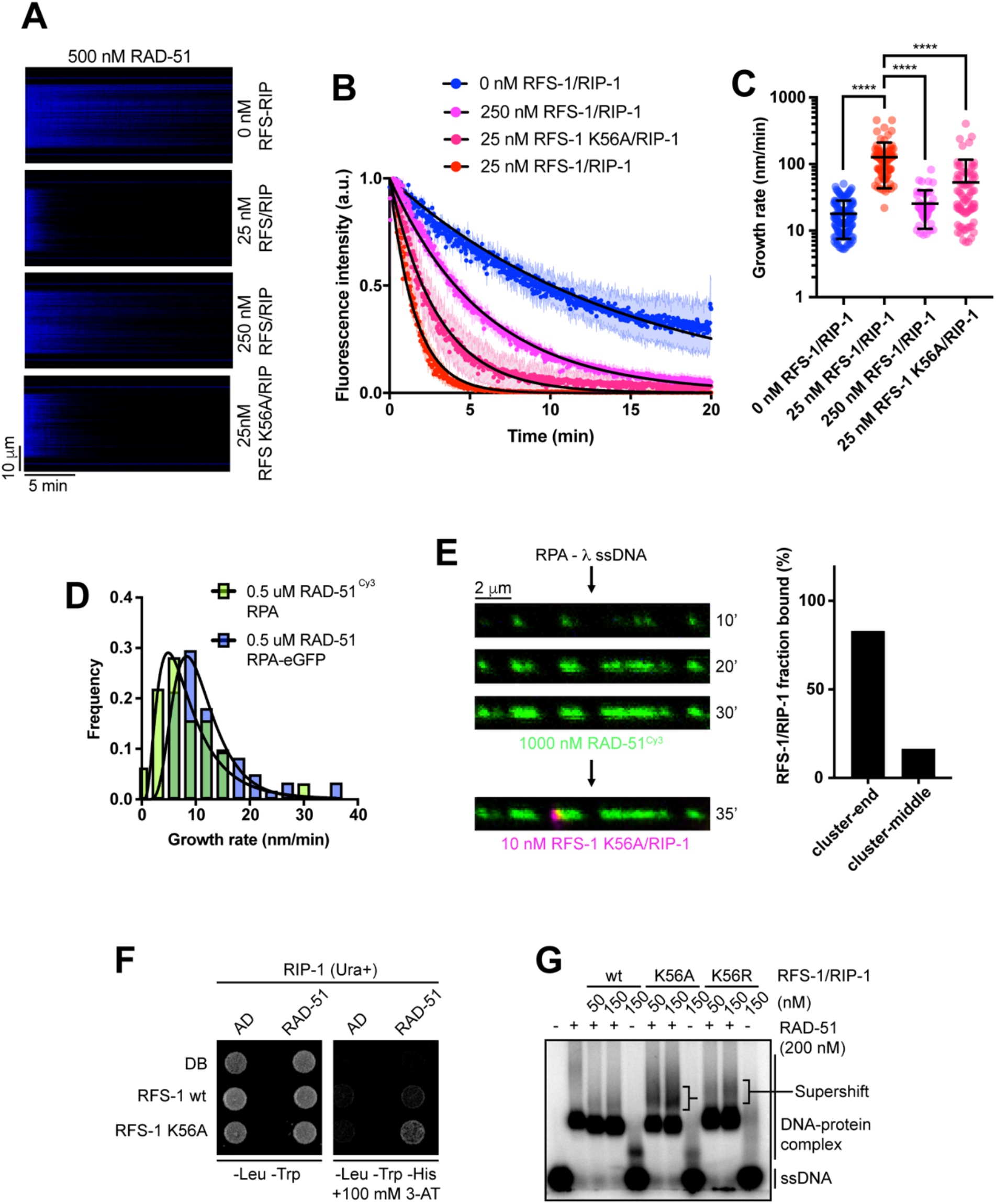
Extended analysis of RFS-1/RIP-1 Walker A mutants. **(A)** Kymograph showing the displacement of RPA-eGFP by RAD-51 in indicated conditions. **(B)** Normalized fluorescence intensity for RPA–eGFP signal in indicated conditions; shaded area represents SEM. (n = 3-7). Black lines represent exponential fits. **(C)** Quantification of growth rates in indicated conditions. P < 0.0001. Mann-Whitney test. **(D)** Growth rate distribution of unlabelled RAD-51 on RPA-eGFP coated ssDNA (n = 61; geometric mean of 10.1 ± 1.6 nm/min, 3 independent molecules) and growth rate distribution of RAD-51^Cy3^ on RPA coated ssDNA (n = 32, geometric mean of 7.5 ± 1.9 nm/min, 5 independent molecules). Lines represent lognormal fits. **(E)** Confocal fluorescence images taken after subsequent incubation-detection cycles using RPA-λ ssDNA and 1000 nM RAD-51^Cy3^ (green) or RFS-1 K56A/RIP-1(A647) (magenta). Cumulative incubation time is indicated. Representative images are shown. Quantification of filament end binding frequencies of RFS-1 K56A/RIP-1(A647) from two independent experiments. **(F)** DB-RFS-1 and AD-RAD-51 interaction in yeast two-hybrid, indicated by survival on media containing 3-aminotriazole (3-AT) in the absence of histidine. Growth on media lacking leucine and tryptophan is a positive control for plasmid transfection. RIP-1 was co-expressed in the same strains on a Ura-selectable plasmid. **(G)** EMSA showing formation of additional ssDNA-protein complexes (supershift) when RFS-1 K56A or RFS-1 K56R/RIP-1 variants were used. For these experiments, proteins were mixed together and incubated on ice for 5 min, 1 nM (in molecules) 60-mer radio-labelled ssDNA was added and incubated for 10 min at 25 °C with indicated proteins followed by 10 min crosslinking with 0.25% glutaraldehyde at 25 °C. Protein-DNA complexes were resolved using 1 % agarose gel electrophoresis at 4 °C for 2 h 20 min.

## Notes

### Competing Interest Statement

The authors have declared no competing interest.

## REFERENCES

Amitani, I., Liu, B., Dombrowski, C.C., Baskin, R.J., and Kowalczykowski, S.C. (2010). Watching individual proteins acting on single molecules of DNA. Methods Enzymol 472, 261–291.

Autour, A., S, C.Y.J. A D.C., Abdolahzadeh, A., Galli, A., Panchapakesan, S.S.S., Rueda, D., Ryckelynck, M., and Unrau, P.J. (2018). Fluorogenic RNA Mango aptamers for imaging small non-coding RNAs in mammalian cells. Nat Commun 9, 656.

Bell, J.C., Plank, J.L., Dombrowski, C.C., and Kowalczykowski, S.C. (2012). Direct imaging of RecA nucleation and growth on single molecules of SSB-coated ssDNA. Nature 491, 274–278.

Buisson, R., Dion-Cote, A.M., Coulombe, Y., Launay, H., Cai, H., Stasiak, A.Z., Stasiak, A., Xia, B., and Masson, J.Y. (2010). Cooperation of breast cancer proteins PALB2 and piccolo BRCA2 in stimulating homologous recombination. Nat Struct Mol Biol 17, 1247–1254.

Candelli, A. (2013). Physical Mechanisms of DNA repair: A single molecule perspective. In Department of Physics and Astronomy (Vrije Universiteit Amsterdam: Vrije Universiteit Amsterdam), pp. 188.

Candelli, A., Hoekstra, T.P., Farge, G., Gross, P., Peterman, E.J., and Wuite, G.J. (2013). A toolbox for generating single-stranded DNA in optical tweezers experiments. Biopolymers 99, 611–620.

Candelli, A., Holthausen, J.T., Depken, M., Brouwer, I., Franker, M.A., Marchetti, M., Heller, I., Bernard, S., Garcin, E.B., Modesti, M., et al. (2014). Visualization and quantification of nascent RAD51 filament formation at single-monomer resolution. Proc Natl Acad Sci U S A 111, 15090–15095.

Chapman, J.R., Taylor, M.R., and Boulton, S.J. (2012). Playing the end game: DNA double-strand break repair pathway choice. Mol Cell 47, 497–510.

Chi, P., Van Komen, S., Sehorn, M.G., Sigurdsson, S., and Sung, P. (2006). Roles of ATP binding and ATP hydrolysis in human Rad51 recombinase function. DNA Repair (Amst) 5, 381–391.

Chun, J., Buechelmaier, E.S., and Powell, S.N. (2013). Rad51 paralog complexes BCDX2 and CX3 act at different stages in the BRCA1-BRCA2-dependent homologous recombination pathway. Mol Cell Biol 33, 387–395.

Chung, W.H., Zhu, Z., Papusha, A., Malkova, A., and Ira, G. (2010). Defective resection at DNA double-strand breaks leads to de novo telomere formation and enhances gene targeting. PLoS Genet 6, e1000948.

Galletto, R., Amitani, I., Baskin, R.J., and Kowalczykowski, S.C. (2006). Direct observation of individual RecA filaments assembling on single DNA molecules. Nature 443, 875–878.

Gataulin, D.V., Carey, J.N., Li, J., Shah, P., Grubb, J.T., and Bishop, D.K. (2018). The ATPase activity of E. coli RecA prevents accumulation of toxic complexes formed by erroneous binding to undamaged double stranded DNA. Nucleic Acids Res 46, 9510–9523.

Gutierrez-Escribano, P., Newton, M.D., Llauro, A., Huber, J., Tanasie, L., Davy, J., Aly, I., Aramayo, R., Montoya, A., Kramer, H., et al. (2019). A conserved ATP- and Scc2/4-dependent activity for cohesin in tethering DNA molecules. Sci Adv 5, eaay6804.

Haas, K.T., Lee, M., Esposito, A., and Venkitaraman, A.R. (2018). Single-molecule localization microscopy reveals molecular transactions during RAD51 filament assembly at cellular DNA damage sites. Nucleic Acids Res 46, 2398–2416.

Hashimoto, Y., Ray Chaudhuri, A., Lopes, M., and Costanzo, V. (2010). Rad51 protects nascent DNA from Mre11-dependent degradation and promotes continuous DNA synthesis. Nat Struct Mol Biol 17, 1305–1311.

Hegner, M., Smith, S.B., and Bustamante, C. (1999). Polymerization and mechanical properties of single RecA-DNA filaments. Proc Natl Acad Sci U S A 96, 10109–10114.

Hilario, J., Amitani, I., Baskin, R.J., and Kowalczykowski, S.C. (2009). Direct imaging of human Rad51 nucleoprotein dynamics on individual DNA molecules. Proc Natl Acad Sci U S A 106, 361–368.

Jensen, R.B., Carreira, A., and Kowalczykowski, S.C. (2010). Purified human BRCA2 stimulates RAD51-mediated recombination. Nature 467, 678–683.

Jensen, R.B., Ozes, A., Kim, T., Estep, A., and Kowalczykowski, S.C. (2013). BRCA2 is epistatic to the RAD51 paralogs in response to DNA damage. DNA Repair (Amst) 12, 306–311.

Joo, C., McKinney, S.A., Nakamura, M., Rasnik, I., Myong, S., and Ha, T. (2006). Real-time observation of RecA filament dynamics with single monomer resolution. Cell 126, 515–527.

Kim, D.H., Lee, K.H., Kim, J.H., Ryu, G.H., Bae, S.H., Lee, B.C., Moon, K.Y., Byun, S.M., Koo, H.S., and Seo, Y.S. (2005). Enzymatic properties of the Caenorhabditis elegans Dna2 endonuclease/helicase and a species-specific interaction between RPA and Dna2. Nucleic Acids Res 33, 1372–1383.

King, G.A., Burla, F., Peterman, E.J.G., and Wuite, G.J.L. (2019). Supercoiling DNA optically. Proc Natl Acad Sci U S A.

King, M.C., Marks, J.H., Mandell, J.B., and New York Breast Cancer Study, G. (2003). Breast and ovarian cancer risks due to inherited mutations in BRCA1 and BRCA2. Science 302, 643–646.

Lee, J.Y., Qi, Z., and Greene, E.C. (2016). ATP hydrolysis Promotes Duplex DNA Release by the RecA Presynaptic Complex. J Biol Chem 291, 22218–22230.

Lim, C.J., Zaug, A.J., Kim, H.J., and Cech, T.R. (2017). Reconstitution of human shelterin complexes reveals unexpected stoichiometry and dual pathways to enhance telomerase processivity. Nat Commun 8, 1075.

Lisby, M., Barlow, J.H., Burgess, R.C., and Rothstein, R. (2004). Choreography of the DNA damage response: spatiotemporal relationships among checkpoint and repair proteins. Cell 118, 699–713.

Liu, J., Doty, T., Gibson, B., and Heyer, W.D. (2010). Human BRCA2 protein promotes RAD51 filament formation on RPA-covered single-stranded DNA. Nat Struct Mol Biol 17, 1260–1262.

Liu, J., Renault, L., Veaute, X., Fabre, F., Stahlberg, H., and Heyer, W.D. (2011). Rad51 paralogues Rad55-Rad57 balance the antirecombinase Srs2 in Rad51 filament formation. Nature 479, 245–248.

Loveday, C., Turnbull, C., Ramsay, E., Hughes, D., Ruark, E., Frankum, J.R., Bowden, G., Kalmyrzaev, B., Warren-Perry, M., Snape, K., et al. (2011). Germline mutations in RAD51D confer susceptibility to ovarian cancer. Nat Genet 43, 879–882.

Martin, J.S., Winkelmann, N., Petalcorin, M.I., McIlwraith, M.J., and Boulton, S.J. (2005). RAD-51- dependent and -independent roles of a Caenorhabditis elegans BRCA2-related protein during DNA double-strand break repair. Mol Cell Biol 25, 3127–3139.

Meindl, A., Hellebrand, H., Wiek, C., Erven, V., Wappenschmidt, B., Niederacher, D., Freund, M., Lichtner, P., Hartmann, L., Schaal, H., et al. (2010). Germline mutations in breast and ovarian cancer pedigrees establish RAD51C as a human cancer susceptibility gene. Nat Genet 42, 410–414.

Newton, M.D., Taylor, B.J., Driessen, R.P.C., Roos, L., Cvetesic, N., Allyjaun, S., Lenhard, B., Cuomo, M.E., and Rueda, D.S. (2019). DNA stretching induces Cas9 off-target activity. Nat Struct Mol Biol 26, 185–192.

Paix, A., Schmidt, H., and Seydoux, G. (2016). Cas9-assisted recombineering in C. elegans: genome editing using in vivo assembly of linear DNAs. Nucleic Acids Res 44, e128.

Pellegrini, L., Yu, D.S., Lo, T., Anand, S., Lee, M., Blundell, T.L., and Venkitaraman, A.R. (2002). Insights into DNA recombination from the structure of a RAD51-BRCA2 complex. Nature 420, 287–293.

Petalcorin, M.I., Galkin, V.E., Yu, X., Egelman, E.H., and Boulton, S.J. (2007). Stabilization of RAD- 51-DNA filaments via an interaction domain in Caenorhabditis elegans BRCA2. Proc Natl Acad Sci U S A 104, 8299–8304.

Petalcorin, M.I., Sandall, J., Wigley, D.B., and Boulton, S.J. (2006). CeBRC-2 stimulates D-loop formation by RAD-51 and promotes DNA single-strand annealing. J Mol Biol 361, 231–242.

Raschle, M., Smeenk, G., Hansen, R.K., Temu, T., Oka, Y., Hein, M.Y., Nagaraj, N., Long, D.T., Walter, J.C., Hofmann, K., et al. (2015). DNA repair. Proteomics reveals dynamic assembly of repair complexes during bypass of DNA cross-links. Science 348, 1253671.

Robertson, R.B., Moses, D.N., Kwon, Y., Chan, P., Zhao, W., Chi, P., Klein, H., Sung, P., and Greene, E.C. (2009). Visualizing the disassembly of S. cerevisiae Rad51 nucleoprotein filaments. J Mol Biol 388, 703–720.

Schlacher, K., Christ, N., Siaud, N., Egashira, A., Wu, H., and Jasin, M. (2011). Double-strand break repair-independent role for BRCA2 in blocking stalled replication fork degradation by MRE11. Cell 145, 529–542.

Shahid, T., Soroka, J., Kong, E., Malivert, L., McIlwraith, M.J., Pape, T., West, S.C., and Zhang, X. (2014). Structure and mechanism of action of the BRCA2 breast cancer tumor suppressor. Nat Struct Mol Biol 21, 962–968.

Sigurdsson, S., Van Komen, S., Bussen, W., Schild, D., Albala, J.S., and Sung, P. (2001). Mediator function of the human Rad51B-Rad51C complex in Rad51/RPA-catalyzed DNA strand exchange. Genes Dev 15, 3308–3318.

Somyajit, K., Saxena, S., Babu, S., Mishra, A., and Nagaraju, G. (2015). Mammalian RAD51 paralogs protect nascent DNA at stalled forks and mediate replication restart. Nucleic Acids Res 43, 9835–9855.

Stark, J.M., Pierce, A.J., Oh, J., Pastink, A., and Jasin, M. (2004). Genetic steps of mammalian homologous repair with distinct mutagenic consequences. Mol Cell Biol 24, 9305–9316.

Sternberg, S.H., Redding, S., Jinek, M., Greene, E.C., and Doudna, J.A. (2014). DNA interrogation by the CRISPR RNA-guided endonuclease Cas9. Nature 507, 62–67.

Symington, L.S. (2016). Mechanism and regulation of DNA end resection in eukaryotes. Crit Rev Biochem Mol Biol 51, 195–212.

Taylor, M.R.G., Spirek, M., Chaurasiya, K.R., Ward, J.D., Carzaniga, R., Yu, X., Egelman, E.H., Collinson, L.M., Rueda, D., Krejci, L., et al. (2015). Rad51 Paralogs Remodel Pre-synaptic Rad51 Filaments to Stimulate Homologous Recombination. Cell 162, 271–286.

Taylor, M.R.G., Spirek, M., Jian Ma, C., Carzaniga, R., Takaki, T., Collinson, L.M., Greene, E.C., Krejci, L., and Boulton, S.J. (2016). A Polar and Nucleotide-Dependent Mechanism of Action for RAD51 Paralogs in RAD51 Filament Remodeling. Mol Cell 64, 926–939.

Taylor, M.R.G., and Yeeles, J.T.P. (2018). The Initial Response of a Eukaryotic Replisome to DNA Damage. Mol Cell 70, 1067–1080 e1012.

Thorslund, T., McIlwraith, M.J., Compton, S.A., Lekomtsev, S., Petronczki, M., Griffith, J.D., and West, S.C. (2010). The breast cancer tumor suppressor BRCA2 promotes the specific targeting of RAD51 to single-stranded DNA. Nat Struct Mol Biol 17, 1263–1265.

van Mameren, J., Modesti, M., Kanaar, R., Wyman, C., Peterman, E.J., and Wuite, G.J. (2009). Counting RAD51 proteins disassembling from nucleoprotein filaments under tension. Nature 457, 745–748.

Ward, J.D., Barber, L.J., Petalcorin, M.I., Yanowitz, J., and Boulton, S.J. (2007). Replication blocking lesions present a unique substrate for homologous recombination. EMBO J 26, 3384–3396.

Watkins, L.P., and Yang, H. (2005). Detection of intensity change points in time-resolved single-molecule measurements. J Phys Chem B 109, 617–628.

Xu, J., Zhao, L., Xu, Y., Zhao, W., Sung, P., and Wang, H.W. (2017). Cryo-EM structures of human RAD51 recombinase filaments during catalysis of DNA-strand exchange. Nat Struct Mol Biol 24, 40–46.

Yin, J., Lin, A.J., Golan, D.E., and Walsh, C.T. (2006). Site-specific protein labeling by Sfp phosphopantetheinyl transferase. Nat Protoc 1, 280–285.

Zhang, L., Ward, J.D., Cheng, Z., and Dernburg, A.F. (2015). The auxin-inducible degradation (AID) system enables versatile conditional protein depletion in C. elegans. Development 142, 4374–4384.

Zhao, W., Vaithiyalingam, S., San Filippo, J., Maranon, D.G., Jimenez-Sainz, J., Fontenay, G.V., Kwon, Y., Leung, S.G., Lu, L., Jensen, R.B., et al. (2015). Promotion of BRCA2-Dependent Homologous Recombination by DSS1 via RPA Targeting and DNA Mimicry. Mol Cell 59, 176–187.

